# Extending the seasons at both ends? Understanding the physiological and genetic context required for stay green mediated yield increase in wheat (*Triticum aestivum*)

**DOI:** 10.64898/2026.05.22.727135

**Authors:** Elizabeth A. Chapman, Simon Orford, Rebecca Beeby, Jacob Lage, Simon Griffiths

## Abstract

Flowering time and monocarpic senescence are tightly environmentally and genetically controlled. Typically, ‘early’ flowering and staygreen traits are associated with opposing life-history strategies; stress avoidance versus adaptation; with flowering time an overarching regulator of crop cycle length. We developed RIL populations segregating for *Ppd-1* and *NAM-1* variation, which are otherwise isogenic. Multi-year field experiments enabled exploration and uncoupling of the relationship between heading and staygreen traits. Heading date manipulation enabled introduction of staygreen traits to their target breeding environments, characterised by a ‘hot-finish’. Under moderate stress, we report a 2.9% and 1.9% increase in grain width (*P*<0.0001), and 5.8% and 3.7% increase in TGW (*P*<0.0001), plus significantly greater yield (*P*<0.1) for ‘late’ heading staygreen RILs homozygous for *NAM-A1,* and *NAM-D1* missense variants, respectively. Grain yield increases were proportionate to the delay in senescence, being greater for the *NAM-A1* than the *NAM-D1* variant. For RIL populations segregating for both traits, senescence variation was observed relative to heading-date. Regarding grain yield, the staygreen trait-associated increase in source size could not compensate for the *Ppd-1a* associated pleiotropic reduction in sink size, even under hypothesised continental target breeding environments, with trait competition identified. Therefore, to maximise the benefits associated with staygreen traits, especially in early-heading favouring environments required targeted manipulation of source-sink dynamics, and we propose multiple strategies.

**Highlight:** Staygreen traits were associated with extending grain fill duration, increasing grain width, TGW and grain yield. There appears an antagonist relationship between earlier heading and staygreen traits.

## Introduction

Global wheat (*Triticum* sp.) production spans 215 M ha (CIMMYT, 2017), with wheat second to rice regarding calorific consumption (FAO, 2013). Such success is due to wheat’s adaptability and tolerance to wide ranging and diverse agroclimates (Shewry, 2009). The UK’s maritime climate makes it amongst the highest yielding wheat environments, which averaged 8.8 t ha^-1^ in 2015; a 25-year record high (DEFRA, 2015). With climate change, weather conditions are increasingly heterogeneous, whereupon wheat yields must be sustainably maintained and improved to ensure global food security (Semenov *et al*., 2014).

Final grain yield can be divided into two components, grain number and grain size, with the switch between these determiners occurring over a narrow window (Semenov *et al*., 2014). Grain number determination occurs pre-anthesis, relating to tiller number, spike and spikelet formation, and floret fertility (Brinton and Uauy, 2019; Slafer *et al*., 2023). At anthesis, environmental conditions influence grain number, whereupon frost damage and heat stress reduce spike fertility via sterility or floret abortion, respectively (Cromey *et al*., 1998; Farooq *et al*., 2011).

Pre-and post-anthesis processes influence grain size, being a product of carpel and cell size, cell number, and grain filling rate (Brinton and Uauy, 2019). The palea and lemma physically constrain floret cavity size, illustrating an element of maternal control, with carpel size and final grain weight positively correlated (Brinton and Uauy, 2019), and grain length and width independently genetically controlled (Gegas *et al*., 2010).

Early grain development is characterised by rapid cell proliferation, which sharply declines ∼6 days after anthesis (daa), followed by cell expansion (Brinton and Uauy, 2019). Maximum grain length is determined ∼15 daa, coinciding with the period of maximum grain fill rate, ∼14-28 daa (Neghliz *et al*., 2016; Brinton and Uauy, 2019). At ∼40 daa, maximum dry grain weight is recorded and grain moisture content maintained, before rapid water loss and physiological maturation. Combined, grain weight improvement arises from differences in cell expansion, and, or proliferation, alongside altered sugar metabolism and cell wall properties (Brinton and Uauy, 2019; Slafer *et al*., 2023).

With pre-and post-anthesis events significantly influencing grain yield, correct environmental synchronisation of flowering time is crucial. Flowering too early risks frost damage, leading to sterility, or grain deformation if suffered during grain fill (Cromey *et al*., 1998). Late flowering may curtail grain fill due to terminal heat or drought stress, reducing grain weight (Wardlaw, 2002), or delay grain maturation, risking pre-harvest sprouting (Nishida *et al*., 2012).

In wheat, *Ppd-1* (*Photoperiod-1*) is the major regulator of flowering time, promoting flowering under increasing daylengths (Beales *et al*., 2007; Langer *et al*., 2014). Photoperiod insensitive alleles exist for all three homoeologues, arising from promoter deletions, *Ppd-A1a* and *Ppd-D1a* (Nishida *et al*., 2012; Shaw *et al*., 2012; 2013), and copy number variation, *Ppd-B1a* (Díaz *et al*., 2012; Shaw *et al.,* 2013), respectively. *Ppd-D1a* explains ∼58% of genetic variation in flowering time and *Ppd-B1a* just 3.2% (Langer *et al*., 2014). Combining single, double, or triple *Ppd-1a* variants accelerates flowering by 15-30 days (Shaw *et al*., 2012). Multiple *Ppd-1a* sources and haplotypes exist, with their geographic spread illustrating their importance in wheat adaptation (Kiss *et al*., 2014; Muterko *et al*., 2015).

Whereas the floral transition marks the switch between vegetative and reproductive growth, monocarpic senescence is the terminal stage in wheat development. Senescence involves the remobilisation and reassimilation of resources, including 80% of leaf nitrogen and phosphorus into the grain (Buchanan-Wollaston, 2007). With leaves largely maintained until senescence completion, green canopy and grain fill duration are assumed to be associated, with Pinto *et al*. (2016) and De Souza Luche *et al*. (2017) reporting correlations of *r*≥0.35 (*P*≤0.0001) and *r*=0.7 (*P*≤0.01), respectively.

Our previous grain filling experiments conducted for *NAM-1* homoeologous, ‘staygreen’, mutants report grain filling extensions proportional to their observed delays in senescence (Chapman *et al*., 2021b). Spano *et al*. (2003) report that a 10-day delay in the onset of senescence correlated with a respective 10-12% and 7.5-20% increase in TGW and grain yield for a *Triticum durum* EMS mutant compared to parental cv. Trinakria. However, this relationship is complex, and Xie *et al*. (2016) report negative correlations between grain fill duration and green area loss of *r*=-0.40, and between rate of chlorophyll loss and TGW, *r*=-0.44, *P*<0.01.

Differences in senescence progression, including rate, duration, or onset, characterise staygreen phenotypes (Vijayalakshmi *et al*., 2010; Gregersen *et al*., 2013; Chapman *et al*., 2021a). Functional staygreen traits are associated with maintenance of photosynthetic activity, not altered chlorophyll catabolism (Thomas, 2000; Gregersen *et al*., 2013; Thomas and Ougham, 2014). Staygreen traits have been indirectly selected over the last 50 years in Australia (Kitonyo *et al*., 2017) and Western Europe (Voss-Fels *et al*., 2019). Evaluation of 14 Australian wheat cultivars developed between 1958 to 2011 by Kitonyo *et al*. (2017) reports increasing minimum NDVI scores with year of release, with cv. Heron (released 1958) early and slow to senesce, and cv. Justica CL Plus (released 2011) greener overall but rapidly senescing.

Indirect staygreen trait selection may relate to an association with alternative yield-enhancing or physiological traits, including canopy architecture, high biomass, or resource-use efficiency. Assessing 100 diverse wheat lines, Kumari *et al*. (2007) report positive associations between staygreen traits, canopy temperature depression and grain fill duration. Under drought stress, Christopher *et al*. (2008) found the yield advantage afforded by CIMMYT line SeriM82 (staygreen) over Hartog (non-staygreen) was associated with greater biomass and faster grain filling rate. Comparative modelling of drought tolerant versus susceptible wheat ideotypes under 2050 climate change predictions estimate the yield benefits associated with staygreen traits across Central and Eastern European wheat growing regions at 10-23% (Senapati *et al*., 2019). Senescence regulation is complex, involving the co-ordinated integration of environmental and developmental cues for timely resource remobilisation and reassimilation. Previous studies struggled to uncouple senescence-heading date interactions (Bogard *et al*., 2011; Camargo *et al*., 2016; Xie, Mayes and Sparkes, 2016). Essentially, an earlier flowering plant likely senesces earlier, whilst late flowering plants apparently exhibit staygreen phenotypes due to developmental differences.

Multiple senescence QTL are reported (Vijayalakshmi *et al*., 2010; Bogard *et al*., 2011; Camargo *et al*., 2016; Christopher *et al*., 2016; Xie, Mayes and Sparkes, 2016; Ren *et al*. 2022; Li *et al*. 2023; Ren *et al*. 2023) with many environmentally responsive and unstable across years, except *GPC-1* (*Grain Protein Content-1*) (Uauy *et al*., 2006a; 2006b). The *GPC-1* locus encodes *NAM-1* (*No Apical Meristem*), which belongs to the plant specific NAC transcription factor family (Olsen *et al*., 2005) and regulates senescence, grain protein, zinc, and iron content (Uauy *et al*., 2006). Knock down, or mutagenesis, of *NAM-1* delays wheat senescence by ≤3 weeks (Avni *et al*., 2014; Borrill *et al*., 2015; Chapman *et al*., 2021b), with knockdown, or mutagenesis, of paralog *NAM-2* also delaying senescence (Pearce *et al*., 2014; Borrill *et al*., 2019).

*NAM-1* expression peaks around anthesis, with changes in expression triggering dramatic changes in transcriptional activity and differential gene expression (Cantu *et al*., 2011; Avni *et al*., 2014; Borrill *et al*., 2019; Andleeb *et al*., 2023). For example, early in senescence, genes associated with ion and transmembrane transport, protein metabolism and stress and hormonal responses are upregulated, and those associated with starch biosynthesis, housekeeping roles, defence and photosynthesis downregulated (Cantu *et al*., 2011; Pearce *et al*., 2014; Borrill *et al*., 2019). Here, we explore the agronomic utility of two staygreen traits, which we identified as underpinned by homoeologous *NAM-A1* and *NAM-D1* mutations (Chapman *et al*., 2021b), plus targetted dissection of source-sink relationships using precise germplasm.

Multi-year testing was performed to determine whether the benefits of the observed grain fill extension are realisable under UK conditions, where the wheat growing season is already long (Mueller *et al*., 2015). Through developing RIL populations segregating for heading and senescence traits (*Ppd* x staygreen) we control, and uncouple, the known anthesis-senescence relationship (Bogard *et al*., 2011). *Ppd-1a* allelic introgression enabled staygreen trait introduction into our target breeding environments, facilitating exposure to late season heat stress in France, and drought in Germany.

When coupling ‘early’ heading and staygreen traits we tested multiple hypotheses (Fig. 1). Firstly, we predict synergism between earlier flowering and staygreen traits, whereupon an extended grain fill duration would increase yield. Conversely, early flowering and staygreen traits represent opposing life history strategies, stress avoidance versus tolerance, promoting resource competition. Alternatively, combining the two traits could lead to yield stability. Analysing the results of these contrasting heading-senescence combinations, we critique Semenov *et al*. (2014) who postulated adoption of escape, over adaptative, traits as the easier strategy for development of climate-adaptable wheat ideotypes.

**Figure 1.**
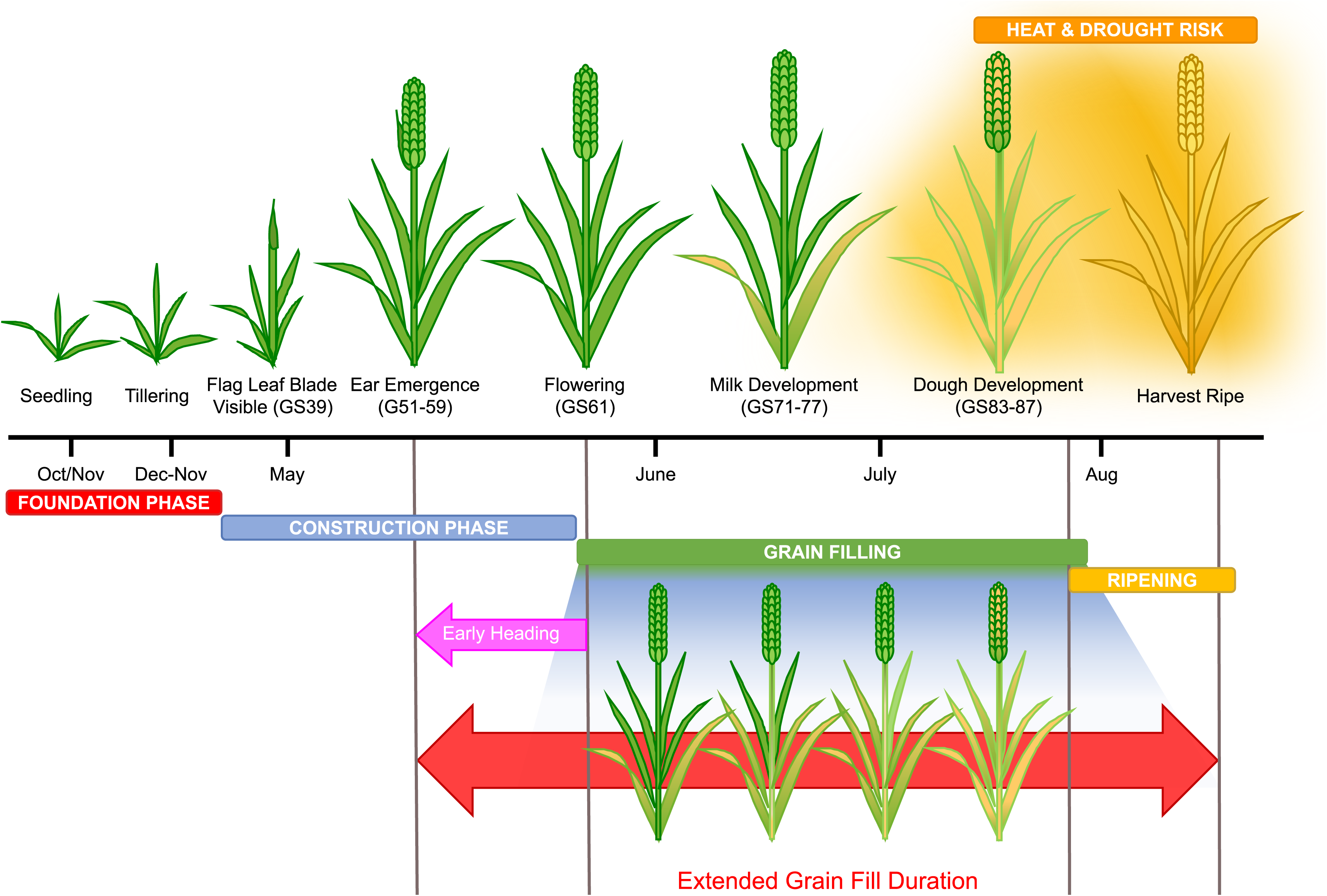
Hypothesized synergism between ‘early’ heading and staygreen traits. Both ‘early’ heading and staygreen traits are associated with longer grain filling duration and increasing TGW, for which trait combining could further enhance grain size and final grain yield. Conversely, earlier heading is typically selected for avoidance of late season stresses that curtail grain filling, whilst staygreen traits are associated with stress adaptation. Thus, combining stress avoidance (heading) and stress adaptive (senescence) traits could result in trait antagonism, penalizing yield.

Through assessing grain yield, yield components, and grain protein content we identify the potential benefits, or penalties, associated with staygreen trait adoption. In response, we report the potential utility of staygreen traits, derived from unexploited novel genetic variation, in breeding, proposing strategies concerning trait adoption.

## Materials & Methods

### Plant Material

Two *Triticum aestivum* cv. Paragon EMS staygreen mutants encoding deleterious missense mutations in *NAM-A1* (T159I; 1189a) and *NAM-D1* (G151E; 2316b) form the basis of this study (Chapman *et al*., 2021b). Two RIL (Recombinant Inbred Line) populations were developed via single seed descent (SSD) for each mutant segregating for senescence (Paragon x staygreen; F_4_), and heading (*Ppd-1* alleles) traits (Ppd x staygreen; F_3_), n=85-96, described previously (Chapman *et al*., 2021b).

## Phenotypic and Agronomic Characterization

### Field Experimentation

#### Paragon x staygreen

In-field assessment of Paragon x staygreen RILs was performed between 2016-2018 at Church Farm (JIC), Norwich (52°38′N, 1°10′E), detailed in Chapman *et al*. (2021a).

#### Ppd x staygreen

Multi-location in-field assessment of *Ppd* x staygreen RILs was performed by KWS between 2016-2018 (Supplementary Table S1), with selections informed by phenotypic data collected during in-field seed multiplication in summer 2015 at Church Farm, Norwich. RILs were classified as ‘early’ or ‘late’ heading (relative to standard UK phenology) based on ear emergence (GS55; Zadoks, et al., 1974), then senescence type (staygreen or non-staygreen) using mean visual leaf senescence score (Chapman *et al*., 2021a). Initially, 9 representative RILs were selected per combination;-‘early’ or ‘late’ heading ± staygreen; totalling 36 per *Ppd* x staygreen population. Subsequently, the number of RILs per trait combination was reduced in favour of greater replication and multi-environment testing (Table 1). In all years, *Ppd* x staygreen RIL subsets were grown as 1 m^2^ 3-row plots, maximum 2 replicates, for observation and seed multiplication at Church Farm, Norwich.

**Table 1.**
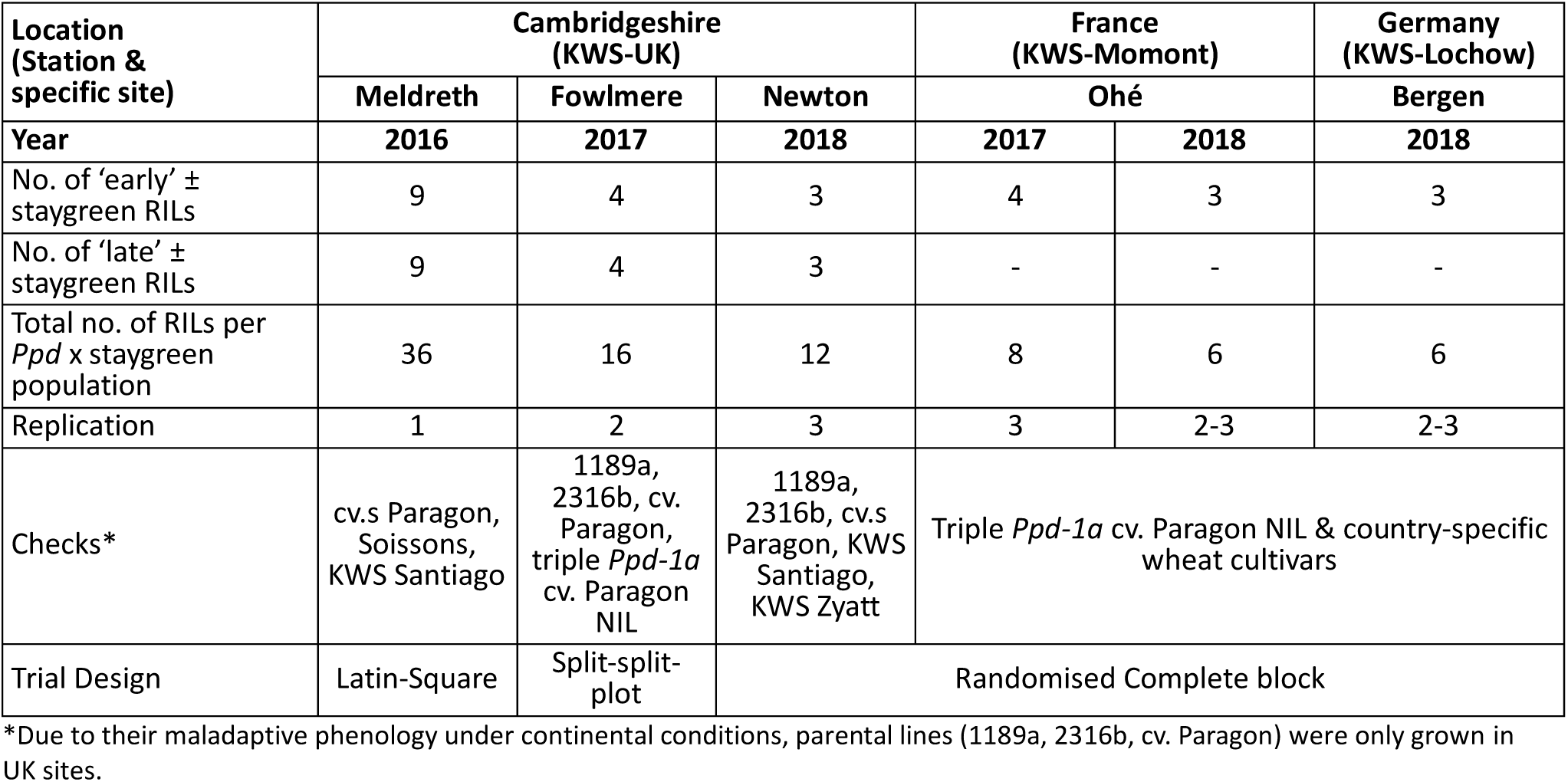
Summary of trial design for each year-site and RILs tested per trait combination.

| Location<br>(Station &<br>specific site) | Cambridgeshire<br>(KWS-UK) |  |  | France<br>(KWS-Momont) |  | Germany<br>(KWS-Lochow) |
| --- | --- | --- | --- | --- | --- | --- |
|  | Meldreth | Fowlmere | Newton | Ohé |  | Bergen |
| Year | 2016 | 2017 | 2018 | 2017 | 2018 | 2018 |
| No. of ‘early’ ± staygreen RILs | 9 | 4 | 3 | 4 | 3 | 3 |
| No. of ‘late’ ± staygreen RILs | 9 | 4 | 3 | - | - | - |
| Total no. of RILs per *Ppd* x staygreen population | 36 | 16 | 12 | 8 | 6 | 6 |
| Replication | 1 | 2 | 3 | 3 | 2-3 | 2-3 |
| Checks* | cv.s Paragon, Soissons, KWS Santiago | 1189a, 2316b, cv. Paragon, triple *Ppd-1a* cv. Paragon NIL | 1189a, 2316b, cv.s Paragon, KWS Santiago, KWS Zyatt | Triple *Ppd-1a* cv. Paragon NIL & country-specific wheat cultivars |  |  |
| Trial Design | Latin-Square | Split-split-plot | Randomised Complete block |  |  |  |

\*Due to their maladaptive phenology under continental conditions, parental lines (1189a, 2316b, cv. Paragon) were only grown in UK sites.

### Experimental Site Descriptions and Treatments

For assessment of *Ppd* x staygreen RILs, site selection was based on location and soil type. In Cambridgeshire, soils consisted of sandy clay loam, with soils of French and German locations described as deep calcareous, and sandy clay, respectively. Drilling rate ranged from 250 to 300 seeds m^-2^ across 8 or 10 rows. Plots measured ≥5 m^2^, and were drilled from mid-late October 2015-2017 in Cambridgeshire, 17^th^ October 2016 and 18^th^ October 2017 in France, and 29^th^ September 2017 in Germany.

Nitrogen application was split across 2-4 applications between March and June. In Cambridgeshire, nitrogen application totalled 204 kg N ha^-1^ in 2016 and 2017, and 149 kg N ha^-1^ in 2018. In France, nitrogen was applied at a rate to achieve a target yield of 8.5 t ha^-1^, equating to 120 kg N ha^-1^ (residual of 126 kg N ha^-1^) and 167 kg N ha^-1^ (residual N of 85 kg N ha^-1^) in 2017 and 2018, respectively. In Germany, nitrogen application totalled 190 kg N ha^-1^. All plots received standard fungicide and herbicide treatment.

### Climatic Data

Supplementary Fig. S1 provides mean daily temperature and rainfall data, sourced from the nearest available weather stations for Church Farm (52°38′N, 1°10′E, Norwich), Royston (52°2′N, 0°1′W, Cambridgeshire; http://dajda.net/), Chartres (48°27’N, 1°30E, France; https://en.tutiempo.net/climate/ws-71430.html) and Bergen (52°48N, 9°55E, Germany: https://opendata.dwd.de/climate_environment/CDC/).

### Phenotypic assessment

Table 2 summarises phenotypes recorded for all year-site combinations. Methodology concerning ear emergence (GS55) (Zadoks *et al.,* 1974), senescence and grain maturity assessment are described in Chapman *et al*. (2021a). In absence of temporal assessment, including 2016, and continental sites, one-off measurements were performed. Plant height was measured for a single, representative, central tiller per plot, excluding awns, to the nearest 1 cm, or 5 cm, in Norwich and Cambridgeshire, and France and Germany respectively.

**Table 2.**
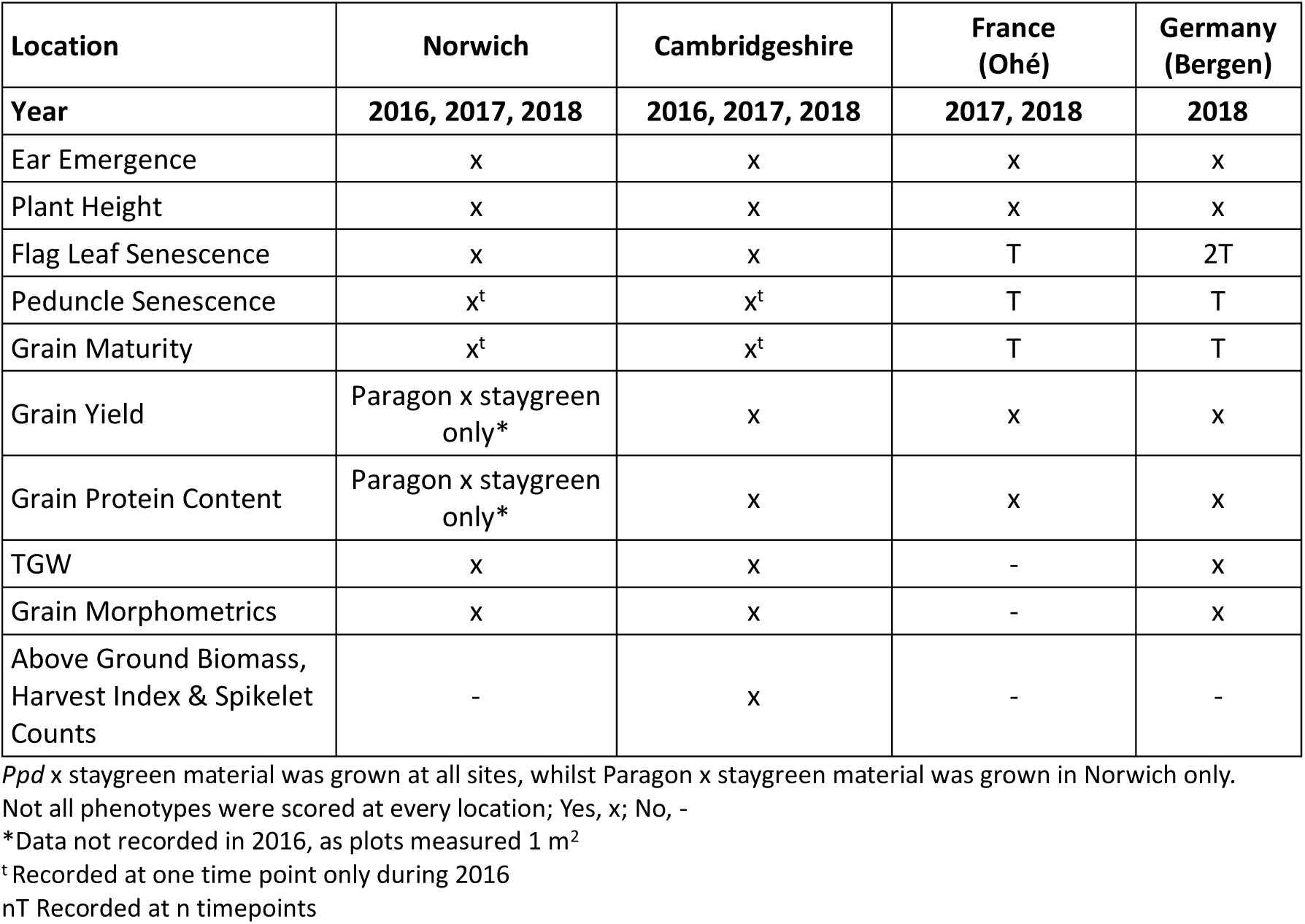
Summary of phenotypes recorded for each year-site combination.

Grain yield was recorded per plot when combine harvested and adjusted according to plot area and grain moisture content. To determine grain protein content Near Infrared Spectroscopy (NIRS) was performed using either flour or grain samples, *Ppd* x staygreen and Paragon x staygreen respectively. For flour samples, 5 g of grain was milled at 14 000 rpm using a Retsch Cyclone mill (Retsch GmbH, Germany), and stored at 8 °C until required. Grain samples weighed ∼20 g, and grain protein content measured using a Perten DA7250 NIR analyser (Perkin Elmer, Waltham, MA). Grain morphometrics, including TGW, width, length, and area were obtained using a MARVIN grain analyser (GTA Sensorik GmbH, Germany), n=300 to 400 grains.

To determine the combined influence of phenology and senescence on grain yield, yield components were assessed for *Ppd* x staygreen trials (UK only). To measure above ground biomass and harvest index (HI), two 30 cm row sections of plants were pulled from the 3^rd^ or 4^th^ row, 30 cm deep into the plot from each end before combine harvesting. Here, roots were left attached, excess soil shaken off, plants gathered in the same orientation and placed, ears first, into a cellophane bag secured using a cable tie around plant stems. Samples were dried for longer term storage prior to subsequent processing, beginning with root removal, with no more than 1-2cm of stems removed, and plants weighed to determine above ground biomass. Ears were removed and threshed using a laboratory thresher (Wintersteiger AG, Austria), chaff checked for seed, and remaining chaff removed. Resulting seeds were weighed and counted using a Seed Count S-25 (Wintersteiger AG, Austria). To calculate HI and TGW, grain weight was divided by above ground biomass, and grain number multiplied by 1000, respectively.

For UK grown *Ppd* x staygreen material, grab sampling was performed to calculate spike yield components. Before harvest, 30 ears were randomly sampled, placed into pre-labelled brown paper bags and dried for longer term storage. Per grab sample, 5 ears were randomly selected, spikelets counted, and ears per bag recounted. Ears were threshed, chaff removed, and seed weighed and counted. As a measure of spike fertility, seed number was divided by the number of ears, and by spikelet number, to calculate grains ear^-1^ and grains spikelet^-1^, respectively.

### Grain Filling Experiments

To test if combining early-heading and staygreen traits further extended grain fill relative to 1189a and 2316b and cv. Paragon (Chapman *et al*., 2021b) grain filling experiments were conducted for RILs *Ppd* x 1189a-56 and *Ppd* x 2316b-57 in summer 2019. These RILs, previously included in all multi-location experiments, carry *Ppd-1a* (early flowering) alleles and are homozygous for their respective *NAM-1* mutations. For comparison, grain filling experiments were conducted for the parental components, triple *Ppd-1a* NILs, cv. Paragon, 1189a and 2316b. To assess grain filling dynamics grain weight and moisture content were recorded from anthesis (GS61: Zadoks *et al*., 1974) to maturation at 3-4 day intervals, as per Chapman *et al*., (2021b).

## Data Analysis

Data analysis was performed using R version 3.6.0 (R Core Team, 2018) within RStudio (RStudio team, 2015) and data manipulated using the packages ‘data.table’ (Dowle *et al*., 2019), ‘dplyr’ (Wickham *et al*., 2018), ‘plyr’ (Wickham, 2015) and ‘tidyr’ (Wickham *et al*., 2019). Agronomic performance was analysed using linear mixed modelling using the packages ‘lme4’ (Bates *et al*., 2019) and ‘lmerTest’ (Kuznetsova *et al*., 2017). Due to significant differences between year-site combinations experiments were analysed separately. For simultaneous assessment of all genotypes and test for specific effect of contrasting *NAM-1* alleles, a nested structure was adopted with RILs nested by population.

When analysing each trial, a complete linear mixed model was initially applied. Fixed effect terms relate to spatial variation (replicate, row, column), and senescence type (*NAM-1* genotype). Experimental genotypes were categorised as ‘control’ or ‘test’ varieties, referring to parental genotypes and check cultivars, or experimental RILs, respectively. For analysis of *Ppd* x staygreen RIL performance, heading class was considered a fixed effect, and interaction tested. *Ppd* x staygreen RILs were classified as ‘early’ or ‘late’ heading relative to parental cv. Paragon based on phenotypic data. Selection of *Ppd* x staygreen RIL subsets prevented inclusion of *Ppd-1* allelic composition of due to too few degrees of freedom. Retention of fixed effect terms was guided by ANOVA, with non-significant terms (*P*>0.05) dropped iteratively.

A single random effect term was included within the model of ‘genotype’ within ‘population’ (1|population:genotype). Term inclusion enables consideration of random variation occurring between closely related RILs, potentially from underlying background mutations, and retained when considered significant (*P*<0.05) following running the command ‘rand(model)’.

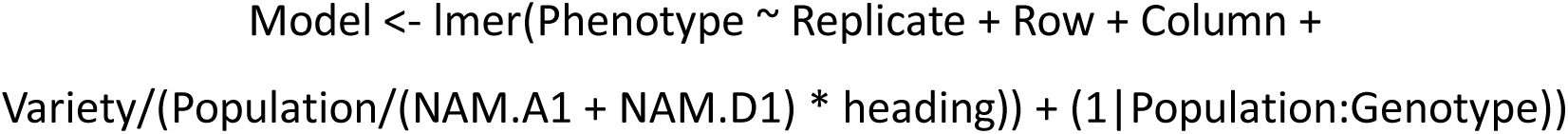

Time-course senescence data and grain filling experiments were analysed as before (Chapman *et al*., 2021b). Models applied were checked for goodness of fit through visual assessment of QQ-plots and that residuals vs. fitted values were normally distributed. To determine significance of observed differences between parents and RIL subsets pairwise Tukey post-hoc tests were conducted using lsmeans() and the ‘list’ option. Graphs were constructed using ‘ggplot2’ (Wickham, Chang, *et al*., 2018), and arranged using R package ‘gridExtra’ (Auguie and Antonov, 2017).

## Results

### Increased grain width associated with NAM-1 variation increases TGW & grain yield of staygreens

Staygreen phenotypes of *Triticum aestivum* cv. Paragon 1189a (*NAM-A1*) and 2316b (*NAM-D1*) mutants are characterised by delays in senescence onset (Chapman *et al*., 2021b). In 2017 and 2018, the UK experienced reduced rainfall and elevated temperatures (Supplementary Fig. S1), curtailing senescence duration by ∼14 and 21 days compared to 2016, respectively. Senescence phenotypes of Paragon x staygreen RILs homozygous for contrasting *NAM-1* mutations were consistently different, *P*<0.05, confirming environmental stability of staygreen phenotypes (Fig. 2; Chapman *et al*., 2021b).

**Figure 2.**
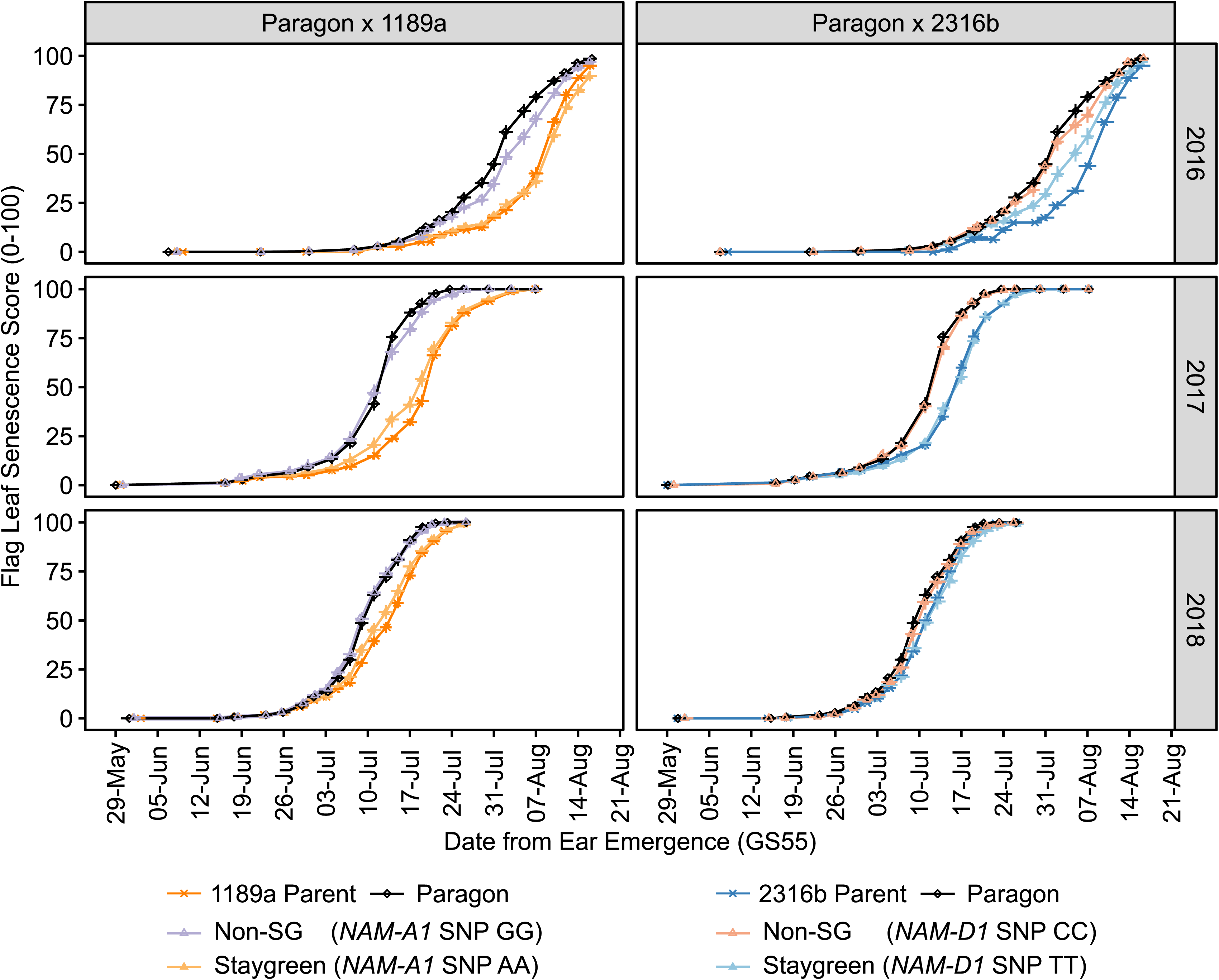
Senescence phenotypes of Paragon x staygreen RIL populations are environmentally stable. Progression of flag leaf senescence of Paragon x 1189a (right) and Paragon x 2316b (left) RILs when grouped by *NAM-1* composition, plus parental lines (Paragon, black; 1189a, dark orange; 2316b, dark blue), Norwich, 2016-2018. ‘Non-staygreen’ (Non-SG; purple, 1189a; red, 2316b) and ‘staygreen’ (orange, 1189a: blue, 2316b) refers to whether RILs are homozygous for the mutant or wildtype allele, respectively. Senescence was scored visually using a 0-100 scale 3-4 times per week from ear emergence (GS55), Mean±SEM, n≥15 per allelic group, n=1-3 per RIL. Supplemental Figure 2 plots senescence against thermal time (°C day).

For 1189a and 2316b, delays in onset of senescence were identified as associated with grain filling extension, potentially enhancing TGW (Chapman *et al*., 2021b). Grain maturity assessment of Paragon x staygreen RIL populations confirmed this, discounting involvement of background mutations (Chapman *et al*., 2021a). We hypothesise this grain filling extension may increase resource delivery to the grain, increasing TGW and final grain yield. To test this, final grain yield and grain morphometric components were recorded for Paragon x staygreen RIL populations grown between 2016-2018 (Fig. 3; Supplementary Table S2).

**Figure 3.**
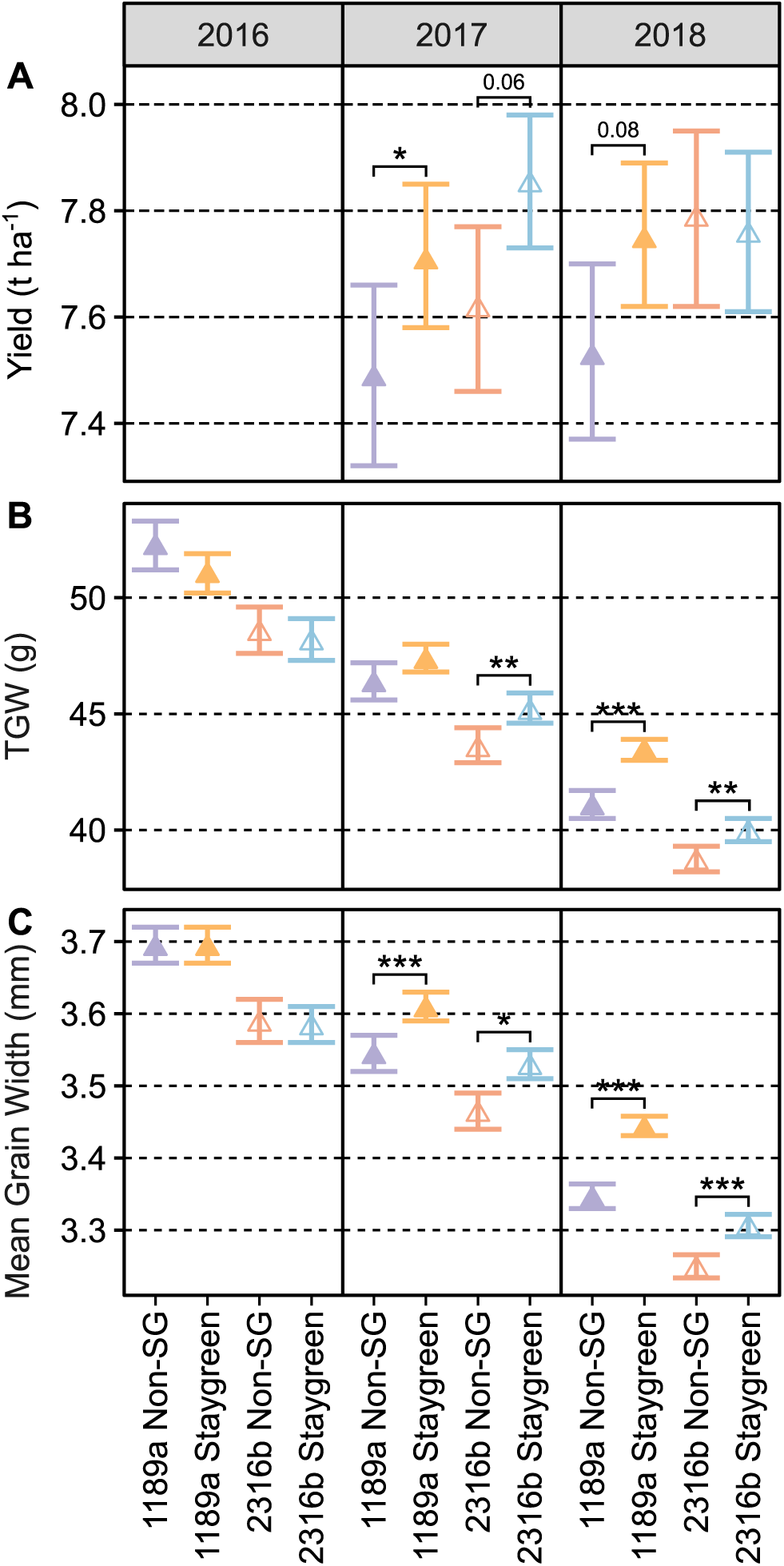
Differences in final grain yield correspond to staygreen-mediated increases in mean grain width & TGW. **(A)** Final grain yield (t ha^-1^), **(B)** TGW (g), **(C)** Mean grain width (mm) of Paragon x staygreen RILs grown between 2016-2018 at Church Farm, Norwich. In 2017 and 2018, mean grain widths of homozygous Paragon x staygreen RILs contrasting for *NAM-1* alleles were significantly different, *P*<0.05, in favour of the staygreen trait (C). The trend for increasing grain width translated into significant increases in TGW (B; *P*<0.001, 1189a (*NAM-A1*), 2018 only; *P*<0.01, 2316b (*NAM-D1*), 2017 & 2018), and final grain yield (A; *P≤*0.08, 1189a (*NAM-A1)*, 2017 & 2018; *P=*0.06, 2316b (*NAM-D1*), 2017 only). Mean±CI_95%_. No differences in grains m^-2^ were observed, *P*≥0.1. RILs grown per population & replication, 2016, n=36, rep.=1; 2017, n=42, rep.=3; 2018, n≥75, rep.=2. TGW (g) and mean grain width (mm) were recorded using the MARVIN grain analyser, sample size n=300-400. ‘Staygreen’ and ‘Non-SG’ refer to RILs homozygous for *NAM-A1* (1189a) or *NAM-D1* (2316b) variants, and cv. Paragon allele respectively. Annotated <0.1, \**P*<0.05, \*\**P*<0.01, \*\*\**P*<0.001 (Tukey *post hoc* test). Grain yield was not recorded in 2016 as experimental plots measured 1 m^2^.

For Paragon x staygreen RILs, *NAM-A1* (1189a) and *NAM-D1* (2316b) variants are associated with increasing specific grain morphometric components, *P≤*0.017, influencing yield (Fig. 3). Final grain yield of Paragon x staygreen RILs homozygous for contrasting *NAM-1* alleles was significantly different upon relaxation of the significance threshold to *P<*0.1 (Fig. 3A), deemed acceptable by plant breeders as such stringency is rarely met. Between 2017 and 2018, Paragon x 1189a RILs homozygous for the *NAM-A1* mutation (staygreen) out-yielded RILs homozygous for the cv.

Paragon allele (non-staygreen) by 2.9%, *P≤*0.076 (Fig. 3A). In 2017, final grain yield of Paragon x 2316b RILs homozygous for the *NAM-D1* mutation was 3.1% greater than those homozygous for the cv. Paragon allele, *P*=0.058, with differences in 2018 not significant, *P=*0.46 (Fig. 3A). In 2016, Paragon x staygreen RILs were grown as 1 m^2^ plots making yield data unreliable.

Contributing to the greater final grain yield of Paragon x staygreen RILs homozygous for *NAM-1* variants is their mean grain width, *P≤*0.02. In 2017, mean grain width of Paragon x staygreen RILs homozygous for either *NAM-1* variant was 1.7% greater compared to those homozygous for the cv. Paragon allele, increasing to +1.9% (2316b, *NAM-D1*) and +2.9% (1189a, *NAM-A1*) in 2018, *P<*0.0001 (Fig. 3C). Grain width is considered a measure of grain filling capacity (Brinton *et al*., 2017), and a determiner of TGW. Between 2017 and 2018, for Paragon x staygreen RILs, the *NAM-D1* variant was associated with a 3.1 to 3.7% increase in TGW, *P≤*0.006, and *NAM-A1* variant a 5.8% increase, *P<*0.0001 (2018 only), Fig. 3B. No differences in grains m^-2^ were observed, *P*≥0.1.

Staygreen phenotypes are source-trait related, likelier influencing fulfilment of grain size potential not grain number (Slafer *et al*., 2023). Regarding grain length (mean, minimum, maximum), and grain width (minimum, maximum), Tukey *post-hoc* tests revealed no differences between homozygous Paragon x staygreen RILs contrasting for *NAM-1* mutations, *P*=0.09 to 1, although in 2016 the *NAM-A1* variant was associated with a 1.7% reduction in mean grain length, *P*=0.02. Results concerning grain area were inconsistent (Supplementary Table S2).

### Uncoupling of flowering and senescence traits

Senescence variation is often explained by heading-date (Camargo *et al*., 2016; Xie, Mayes and Sparkes, 2016), typically correlating with yield and grain protein content (Bogard *et al*., 2011). Phenological manipulation has pleiotropic sink effects, as earlier heading lines reach GS31 sooner, determining final grain number earlier (Zadoks *et al*., 1974; Worland *et al*., 1998; Arjona *et al*., 2018). To control for, identify, and understand heading-senescence interactions, *Ppd* x staygreen RIL populations were developed segregating for both traits in isogenic backgrounds. From each population, ‘early’ and ‘late’ heading ± staygreen RIL subsets were selected and grown in Cambridgeshire, UK, between 2016 and 2018, and yield components recorded (Table 2).

Visual assessment of time-course senescence data of *Ppd* x staygreen RILs illustrates staygreen trait expression in ‘early’ and ‘late’ heading backgrounds (Fig. 4: Supplementary Fig. S2), with differences between heading groups typically significant (*P*<0.05: Supplementary Table S3).

**Figure 4.**
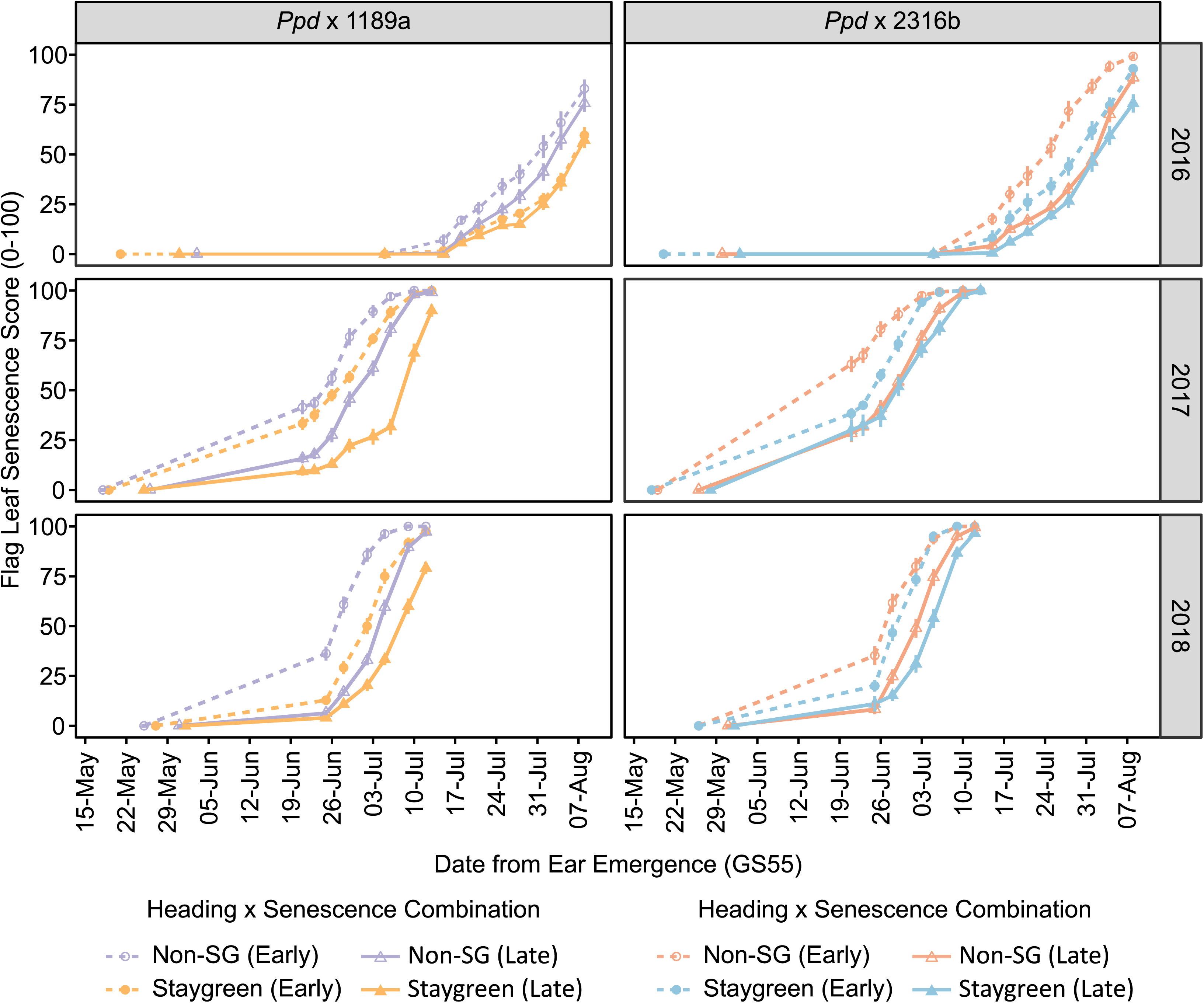
Confirmation of staygreen trait expression for ‘early’ and ‘late’ heading *Ppd* x staygreen RILs. Progression of flag leaf senescence of *Ppd* x 1189a (right) and *Ppd* x 2316b (left) RILs when grouped by heading and *NAM-1* composition, Cambridgeshire, 2016-2018; For Norwich, refer to Supplemental Figure 3. Heading type, ‘early’ (dotted lines), ‘late’ (solid lines). Senescence type, ‘non-staygreen’ (Non-SG; purple, 1189a; red, 2316b), ‘staygreen’ (orange, 1189a; blue, 2316b). ‘Staygreen’ refers to homozygosity for either *NAM-1* variant (*NAM-A1,* 1189a, or *NAM-D1,* 2316b), with the ‘non-staygreen’ group homozygous for the cv. Paragon allele. Senescence was scored visually using a 0-100 scale twice week from ear emergence (GS55), mean ± SEM, n≤15 per heading x senescence combination, n=1-3 per RIL. Supplementary Figure 4 adjusts for heading date variation, plotting senescence against thermal time (°C day).

Differences in senescence of *Ppd* x staygreen RILs homozygous for contrasting *NAM-1* variants, respective and irrespective of heading date, were significant when water limited in 2017 and 2018 (*P*<0.05; Table 3; Supplementary Fig. S1). Through controlling for heading date we successfully uncoupled heading and senescence traits, with effect stronger for *Ppd* x 1189a RILs due to the greater extremity of the *NAM-A1* over the *NAM-D1* (2316b) staygreen phenotype (Table 2; Supplementary Table S3; Avni *et al*., 2014; Chapman *et al*., 2021b).

**Table 3.** NAM-1 mutations influence senescence progression of RILs independent of heading date.

|  |  |  | 2016 |  |  | 2017 |  |  | 2018 |  |  |
| --- | --- | --- | --- | --- | --- | --- | --- | --- | --- | --- | --- |
| RILs | Site | Heading | Ear Emergence<br>(GS55)<br>(dd/mm ± SD) | Staygreen vs. Non-SG<br>P-value |  | Ear Emergence<br>(GS55)<br>(dd/mm ± SD) | Staygreen vs. Non-SG<br>P-value |  | Ear Emergence<br>(GS55)<br>(dd/mm ± SD) | Staygreen vs. Non-SG<br>P-value |  |
|  |  |  |  | Flag Leaf | Peduncle<sup>†</sup> |  | Flag Leaf* | Peduncle |  | Flag Leaf | Peduncle |
| *Ppd* x 1189a<br>(*NAM-A1*) | Cambs. | 'Early' | 21/05 ± 1.8 | 0.002 | - | 18/05 ± 1.3 | 0.03 | 0.08 | 26/05 ± 2.1 | <0.0001 | 0.0005 |
|  |  | 'Late' | 02/06 ± 4.3 | 0.11 | - | 26/05 ± 1.7 | <0.0001 | 0.03 | 31/05 ± 1.2 | <0.0001 | 0.0005 |
|  |  | Overall | - | <0.0001 | - | - | <0.0001 | 0.0003 | - | <0.0001 | 0.0001 |
|  | Norwich | 'Early' | 16/05 ± 3.1 | 0.87 | - | 19/05 ± 2.6 | 0.28 | <0.0001 | 24/05 ± 3.4 | <0.0001 | 0.001 |
|  |  | 'Late' | 30/05 ± 3.7 | 0.08 | - | 28/05 ± 1.8 | 0.0001 | <0.0001 | 01/06 ± 2.0 | <0.0001 | 0.001 |
|  |  | Overall | - | 0.29 | - | - | <0.0001 | <0.001 | - | <0.0001 | 0.0002 |
| *Ppd* x 2316b<br>(*NAM-D1*) | Cambs. | 'Early' | 20/05 ± 1.8 | 0.004 | - | 18/05 ± 1.7 | 0.003 | 0.14 | 26/05 ± 2.2 | 0.06 | 0.22 |
|  |  | 'Late' | 01/06 ± 5.3 | 0.54 | - | 28/05 ± 3.2 | 0.75 | 0.22 | 31/05 ± 2.6 | 0.06 | 0.22 |
|  |  | Overall | - | 0.0008 | - | - | 0.0015 | 0.004 | - | 0.01 | 0.06 |
|  | Norwich | 'Early' | 17/05 ± 4.3 | 0.69 | - | 18/05 ± 2.6 | 0.03 | 0.0012 | 23/05 ± 3.5 | 0.78 | 0.33 |
|  |  | 'Late' | 30/05 ± 6.6 | 0.68 | - | 28/05 ± 3.6 | 0.009 | 0.0012 | 01/06 ± 2.9 | 0.78 | 0.33 |
|  |  | Overall | - | 0.94 | - | - | <0.0001 | 0.0002 | - | 0.35 | 0.09 |

<sup>†</sup>Peduncle senescence not scored in 2016
\*Significant heading date interaction, *P*=0.02 (0.01)

In 2017, the heading-senescence relationship was not entirely escaped. Utilising senescence metrics described in Chapman *et al*. (2021a), significant heading-senescence interactions were reported for TT50, TT70, and TT75 (*P*≤0.04) for *Ppd* x 1189a RILs and TT25 to TT70 (*P*≤0.05) for *Ppd* x 2316b RILs, indicating heading-dependent trait expression. Differences in leaf senescence phenotypes of homozygous *Ppd* x 2316b RILs were significant for ‘early’, not ‘late’, heading RILs, *P≤*0.002 vs. *P>*0.5, respectively, with pattern reversed for *Ppd* x 1189a RILs, *P*≥0.18 vs. *P≤*0.04 (early vs. late) (Supplementary Fig. S2). No significant heading-senescence interactions were reported for peduncle phenotypes, with senescence progression similar in all years (Fig. 4; Supplementary Fig. S2).

### Assessment of grain yield components reveals an interaction between *Ppd-1* and *NAM-1* variation

When coupling ‘early’ heading and staygreen traits we hypothesised a synergistic relationship (Fig. 1). Here, staygreen trait adoption would complement the earlier initiation of grain filling associated with earlier heading via prolonging late season grain fill, delivering additional resources to the grain, increasing final grain yield. However, when manipulating heading date the relationship between senescence and final grain yield is more complex when compared to results for Paragon x staygreen RILs (Fig. 3).

For *Ppd* x staygreen RILs grown in the UK between 2017 and 2018, heading traits had greater influence on final grain yield than senescence (Fig. 5). In 2017, final grain yield of ‘early’ heading *Ppd* x 2316b RILs was 18% greater compared to their ‘late’ heading counterparts, *P=*0.01. In 2018, final grain yields of ‘early’ heading *Ppd* x 1189a and *Ppd* x 2316b RILs were 10.2% and 11.7% lower compared to their respective ‘late’ heading RILs, *P<*0.05. Only in 2018 was a staygreen-trait associated yield increase in final grain yield of 8% reported for *Ppd* x 1189a RILs, *P=*0.06, plus an increasing trend for *Ppd* x 2316b RILs (Fig. 5). In 2016, no differences in final grain yield were reported, *P>*0.05.

**Figure 5:**
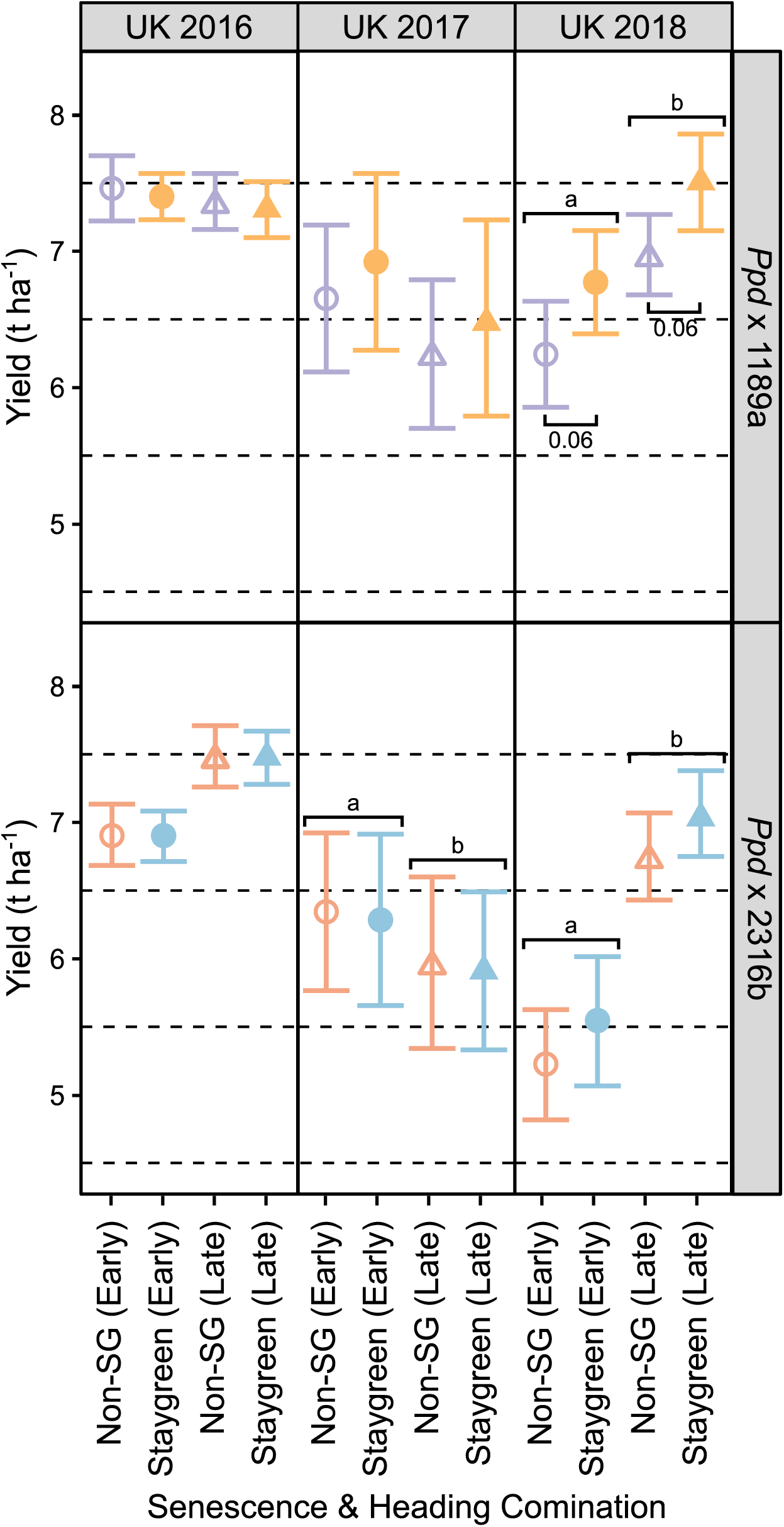
Heading date has opposing effects on final grain yield, whereas staygreen traits appear neutral to positive. Final grain yield (t ha^-1^) for *Ppd* x 1189a (top) and *Ppd* x 2316b (bottom) RILs grouped by heading classification, ‘early’ (circles) or ‘late’ (triangles), Cambridgeshire 2016-2018. Mean ± CI_95%_. ‘Staygreen’ and ‘Non-SG’ refer to RILs homozygous for *NAM-A1* (1189a) or *NAM-D1* (2316b) variants, and cv. Paragon allele respectively. Refer to Supplementary Table S1 for details concerning number of RILs and replicates. ^ab^ Significant differences between heading classes, *P<*0.05. Annotated <,0.1, \**P*<0.05, \*\**P*<0.01, \*\*\**P*<0.001 (Tukey *post hoc* test).

**Figure 6.**
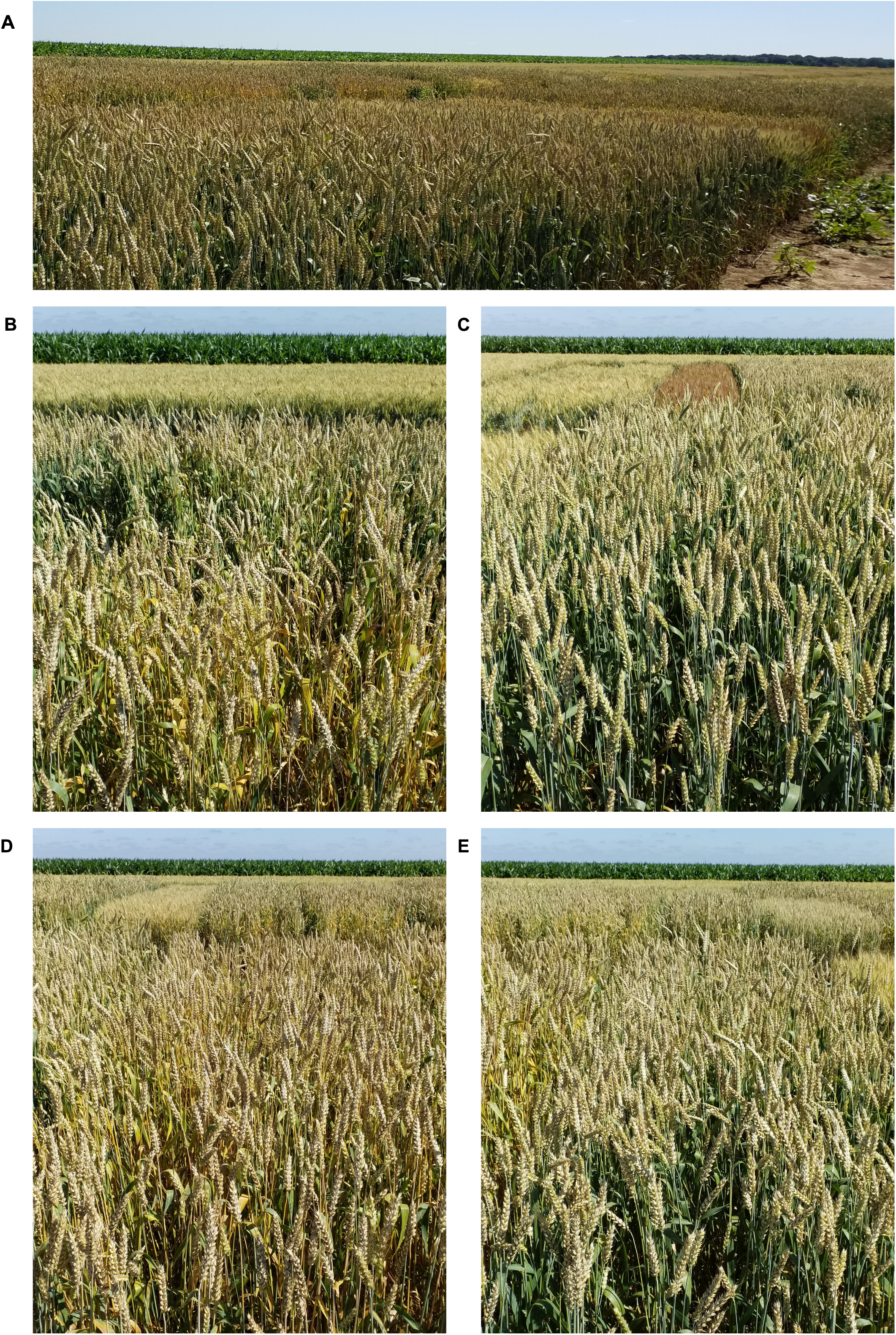
Confirmation of senescence variation for ‘early’ heading *Ppd* x staygreen RILs, France 2018. **(A)** *Ppd* x staygreen trial, Ohé, France, 2018. Senescence of RILs homozygous for the *NAM-A1* (**C**; *Ppd* x 1189a-56), or *NAM-D1* (**E**; *Ppd* x 1189a-16), mutation is delayed relative to those homozygous for cv. Paragon allele (**B**; *Ppd* x 1189a-16), and (**D**; *Ppd* x 2316b-12). Heading date variation was limited to 1-3 days. Photographed 20^th^ June 2018.

**Figure 7.**
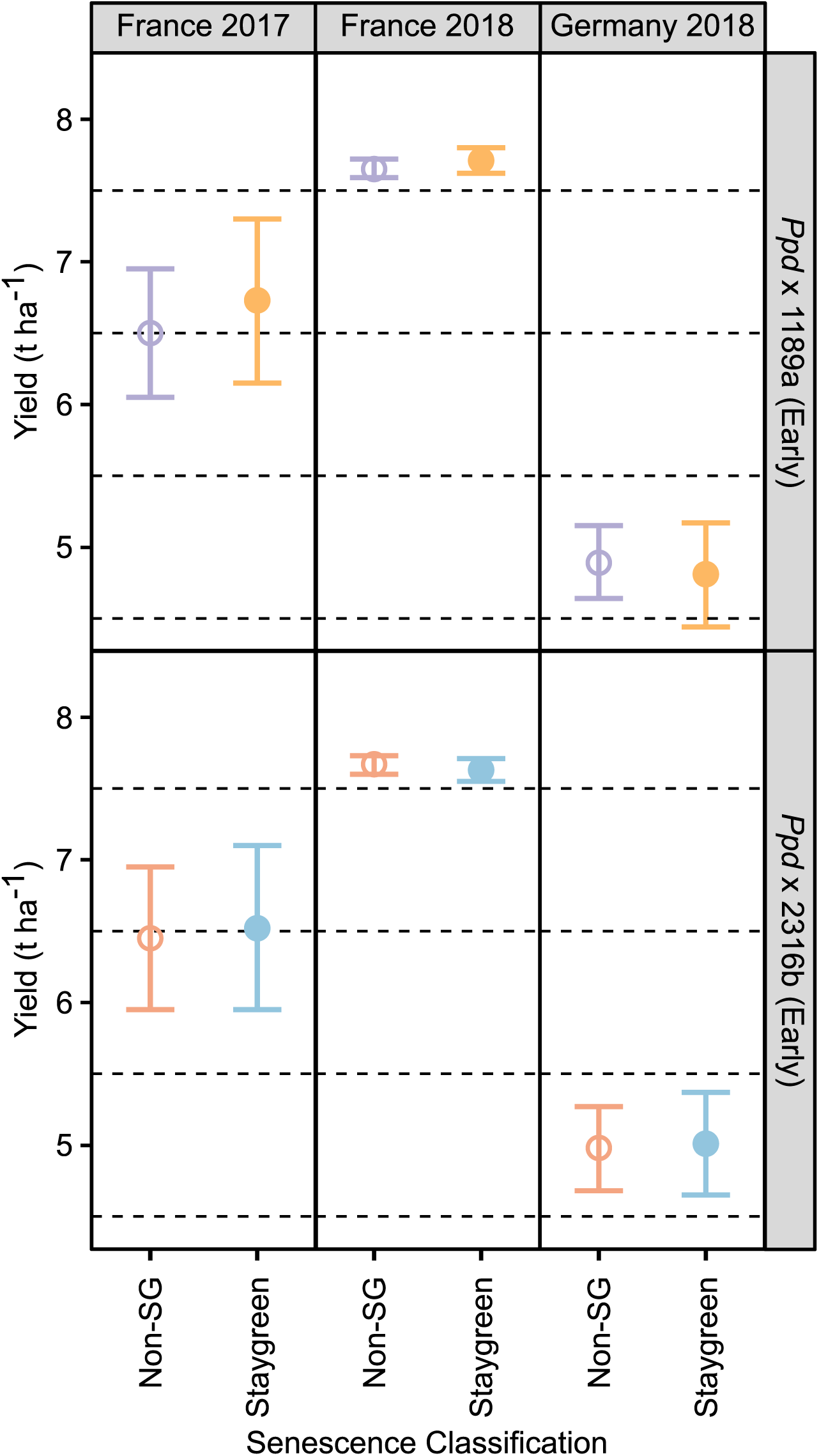
Within our hypothesized target environments the effect of staygreen traits on final grain yield were neutral. Final grain yield (t ha^-1^) of *Ppd* x 1189a (top) and *Ppd* x 2316b (bottom) RILs grown in Ohé, France, 2017 & 2018, and Wohlde, Germany, 2018. Mean±CI_95%_. ‘Staygreen’ and ‘Non-SG’ refer to RILs homozygous for *NAM-A1* (1189a) or *NAM-D1* (2316b) variants, and cv. Paragon allele respectively. Results of Tukey *post-hoc* tests reported no differences in grain yield between senescence types, *P*>0.05. ‘Late’ heading *Ppd* x staygreen RILs were not subjects of multi-location experiments due to their maladaptive phenology. Refer to Supplementary Table S1 for details concerning number of RILs and replicates.

*Ppd-1a* alleles influence multiple grain yield components, specifically grain number, through reducing the period from floral transition to anthesis (Shaw *et al*., 2012; Langer *et al*., 2014). The heading-date dependent differences in grain yield reported for *Ppd* x staygreen RILs concern trade-offs between grain size and grain number. Between 2016 and 2018, the number of spikelets spike^-1^ reported for ‘early’ heading *Ppd* x staygreen RILs was lower compared to ‘late’ heading RILs, *P<*0.05 (excluding *Ppd* x 1189a RILs in 2018). In 2017, this translated into a reduction in grain number m^-2^ between ‘early’ and ‘late’ heading *Ppd* x 1189a and *Ppd* x 2316b RILs of 13.1% and 16.1% respectively, *P≤*0.001. However, these ‘early’ heading *Ppd* x 1189a and *Ppd* x 2316b RILs were higher yielding due to their compensatory 25.1% and 41.1% greater TGWs, respectively (*P≤*0.0001) (Supplementary Table S4). In 2018, the greater TGW of ‘early’ compared to ‘late’ heading *Ppd* x staygreen RILs (*P=*0.015 to 0.11) could not compensate for the associated 15.5 to 19% reduction in grain number, *P<*0.0001, irrespective of senescence (Supplementary Table S4).

Regarding senescence (*NAM-1* allelic composition), no differences were observed for grain number m^-2^, spikelets spike^-1^, grain length (minimum and maximum) for *Ppd* x staygreen RILs (*P*=0.17 to 1). Only in 2018 was the *NAM-A1* variant associated with a 7.3 % increase in the number of spikelets spike^-1^ (P<0.05; Supplementary Table S4). However, in 2018, remobilisation efficiency, according to harvest index, of *Ppd* x 2316b RILs homozygous for the *NAM-D1* mutation was greater, *P=*0.01 (0.17 [0.16, 0.19] vs 0.14 [0.13, 0.16], staygreen vs. non-staygreen; mean ± CI_95%_).

Results for *Ppd* x staygreen RILs and Paragon x staygreen RILs contrast. Here, both homoeologous *NAM-1* variants are associated with increasing mean grain width, TGW and final grain yield (Fig. 3). For *Ppd* x staygreen RILs, heading variation resulted in trade-offs between grain number and grain size, with the contribution of staygreen traits to final grain yield unrealized. To explore this, we analysed grain morphometrics corresponding to multi-location testing of *Ppd* x staygreen RILs (Fig. 8), alongside conducting further grain filling experiments (Fig. 9).

**Figure 8.**
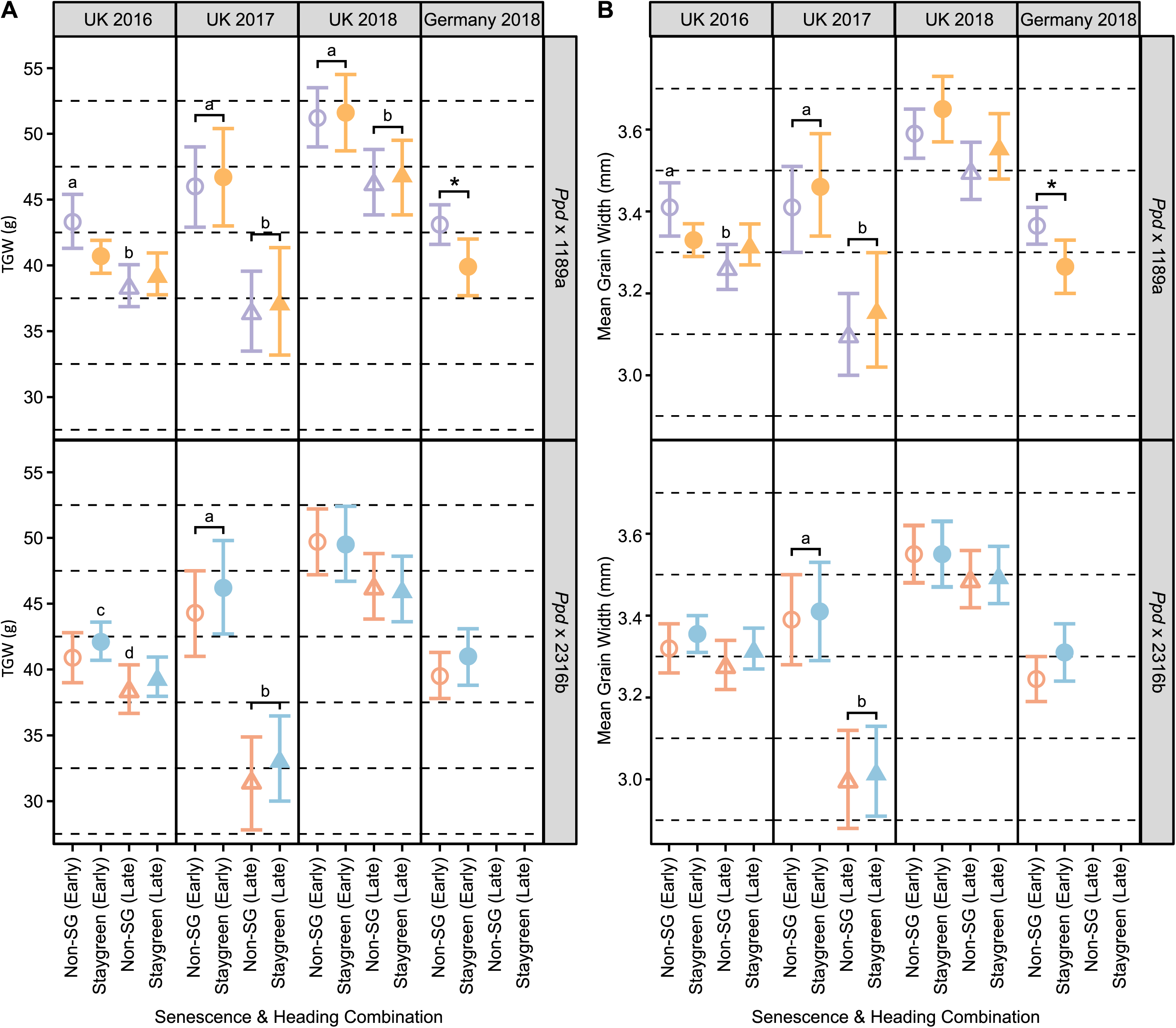
Differences in grain size of *Ppd* x staygreen RILs relate to heading and not senescence variation. **(A)** TGW (g), and **(B)** mean grain width (mm) of *Ppd* x 1189a (top) and *Ppd* x 2316b (bottom) RILs grown in Cambridgeshire, UK, 2016-2018, and Wohlde, Germany, 2018. When both ‘early’ (circles) and ‘late’ (triangles) heading RIL subsets were grown (UK only), grain size of ‘early’ heading RILs was invariably larger. TGW (g) and mean grain width (mm) were recorded using the MARVIN grain analyser, sample size n=300-400. Mean±CI_95%_. ‘Staygreen’ and ‘Non-SG’ refer to RILs homozygous for *NAM-A1* (1189a) or *NAM-D1* (2316b) variants, and cv. Paragon allele respectively. Refer to Supplementary Table S1 for details concerning number of RILs and replicates. \**P*<0.05, ^ab^ Significant differences between heading classes, ^cd^ Significant differences between senescence classifications across heading classes (Tukey *post-hoc* test).

**Figure 9.**
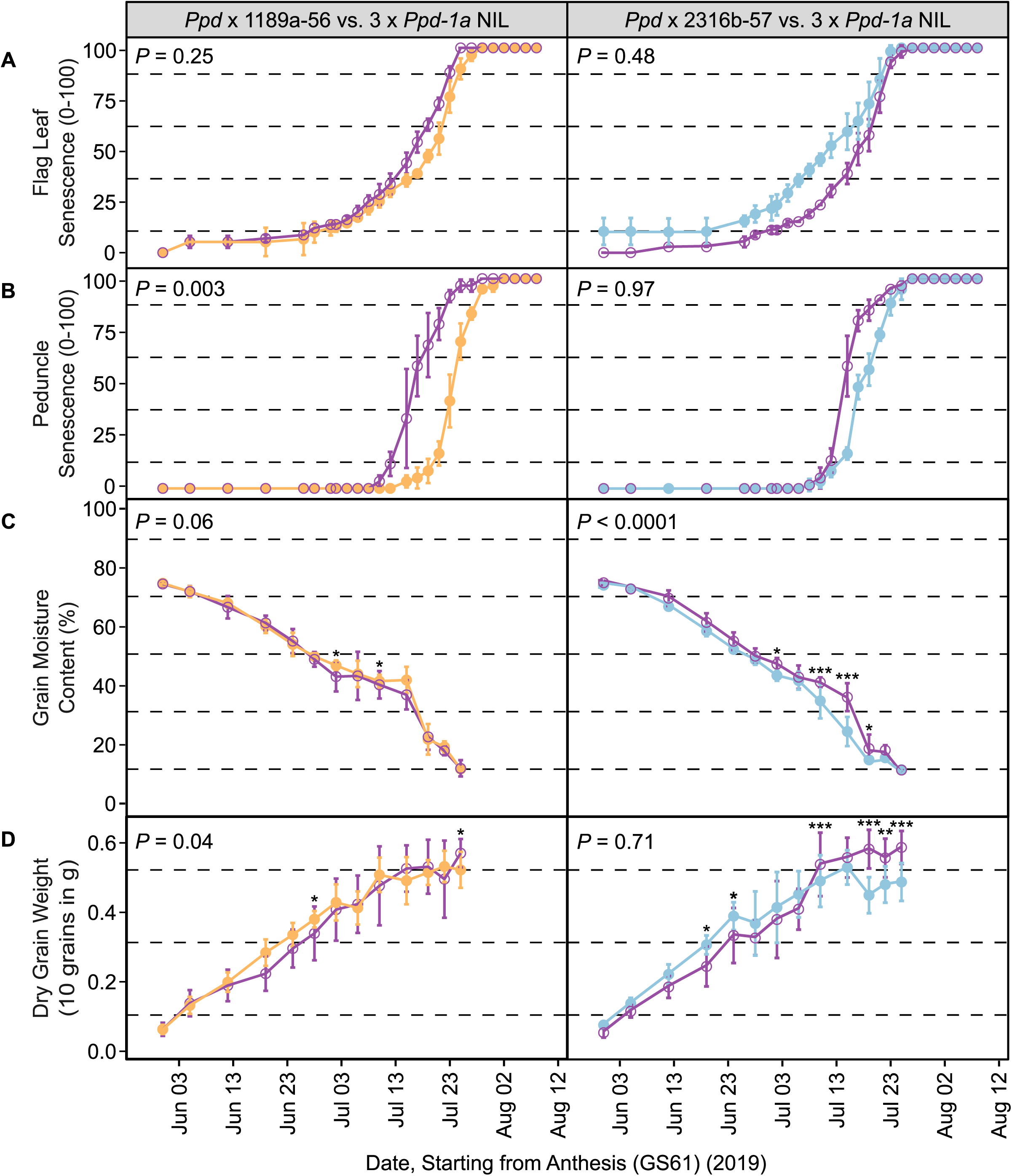
Combining early heading (*Ppd-1a*) & staygreen (1189a, *NAM-*A1; 2316b, *NAM-D1*) traits does not further extend grain fill duration. Grain filling experiments conducted for ‘early’ heading staygreen *Ppd* x 1189a-56 (left, orange) and *Ppd* x 2316b-57 (right, light blue), alongside parental triple *Ppd-1a* cv. Paragon NIL (purple), Norwich, 2019. **(A)** Visual leaf senescence and **(B)** Peduncle senescence; 0-100 scale, mean ± SD. Senescence was scored at the plot level every 2-4 days following anthesis (GS61), n=3. **(C)** Grain moisture content (%) and **(D)** Dry grain weight (10 grains in g), were recorded to determine grain filling dynamics, mean ± SD. 4-5 ears were sampled per plot every 4-6 days from anthesis, 2 plots per genotype. *P*-values represent overall differences (Table 3). Differences at specific time points according to Tukey *post-hoc* tests are indicated (Table 4). *P-*values, *<0.05, **<0.01, ***<0.0001. Confirmation of parental senescence and grain filling phenotypes is provided in Supplemental Figure S5.

**Table 4:**
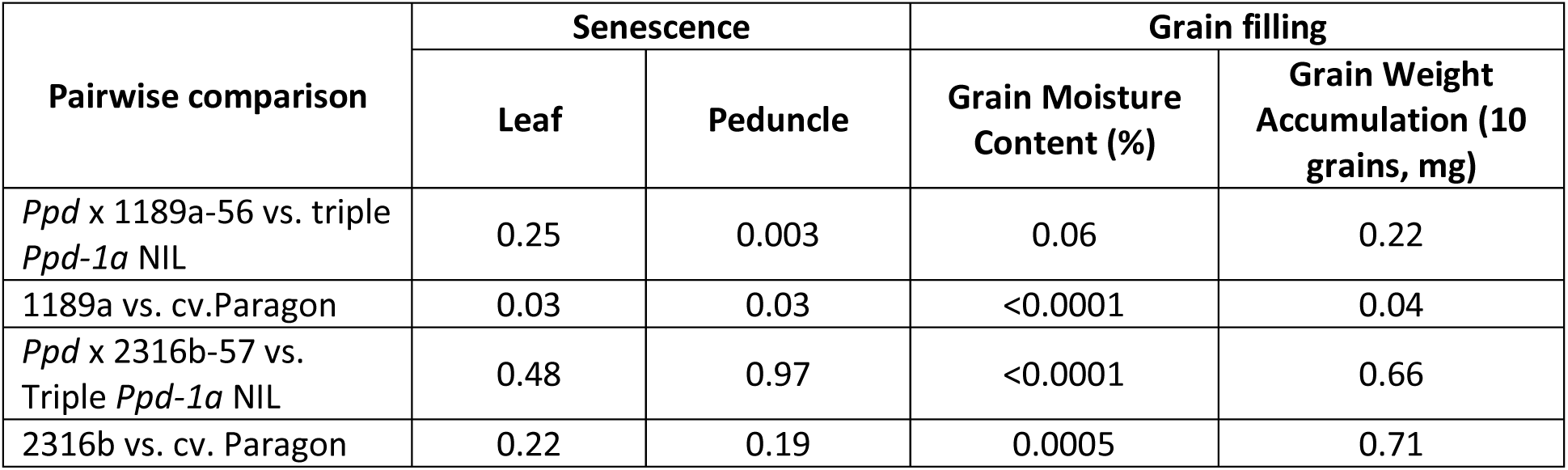
Overall differences in senescence and grain filling phenotypes of ‘early’ ± staygreen & parental lines, 2019.

### Where earlier heading is favoured, *NAM-1* variation delays senescence but does not increase TGW

Phenology of ‘late’ heading Ppd x staygreen RILs is representative of UK adapted wheat cultivars (Worland *et al*., 1998; Langer *et al*., 2014; Senapati *et al*., 2019). However, staygreen traits are associated with conferring environmental stress tolerance (Thomas, 2000; Gregersen *et al*., 2013; Thomas and Ougham, 2014; Jagadish *et al*., 2015), in which *Ppd-1a* alleles are adopted to avoid late season stresses (Fjellheim *et al*., 2014; Langer *et al*., 2014; Muterko *et al*., 2015; Guedira *et al*., 2016). To test the utility of our *NAM-1* variants, and identify their target breeding environment, we grew ‘early’ heading Ppd x staygreen RIL subsets in France & Germany (Table 1; Supplementary Table S1), exposing them to drought and late season stress uncharacteristic of the UK.

In France in 2017 and 2018, mean monthly temperatures for May to August were 1.5 ± 0.4 °C and 1.2 ± 0.5 °C hotter compared to Cambridgeshire, respectively. In 2018, in Germany RILs endured greater water limitation around heading compared to the UK, with May rainfall totalling 19.5 mm versus 45.2 mm, and mean monthly temperature of 16.4 °C versus 14.8 °C (Supplementary Fig. S1). Despite visual expression of staygreen phenotypes under the hypothesised target environments (Fig. 6), no senescence-dependent differences in final grain yield of ‘early’ heading *Ppd* x staygreen RILs were observed, *P≥*0.28 (Fig. 7).

Despite the greater mean monthly average temperatures, *Ppd* x staygreen RILs were exposed to greater water limitation in the UK than France (Supplementary Fig. S1). In Cambridgeshire, June rainfall totalled 47.8 mm and 1.7 mm in 2017 and 2018, respectively, with 56.7 mm and 78.7 mm of rainfall recorded for French sites. In the UK, water limitation identified staygreen traits as conferring a yield advantage, illustrated by the greater final grain yield of Paragon x staygreen RILs homozygous for *NAM-1* variants in 2017, and 2018 (1189a only), *P<*0.1 (Fig. 3).

Greater final grain yields of Paragon x staygreen RILs homozygous for *NAM-1* variants were due to their greater mean grain width, and ultimately TGW (Fig. 3). A similar pattern was not observed for *Ppd* x staygreen RILs (Fig. 8). Analysis of mean grain width and TGW of *Ppd* x 2316b RILs reveals a consistent, albeit insignificant (*P≥*0.3), increasing trend in favour of the *NAM-D1* variant (Fig. 8). For 1189a (*NAM-A1* variant), *Ppd-1a* allelic introduction complicates the positive relationship between staygreen traits and grain size previously reported (Fig. 3; Fig. 8). In the UK in 2016 significant heading-senescence interactions were reported for *Ppd* x 1189a RILs across multiple grain morphometrics, including TGW, mean grain width and maximum grain length, *P≤*0.03 (Supplementary Table S4). This interaction contributed to the differences in TGW, and mean grain width observed between ‘early’ and ‘late’ heading *Ppd* x 1189a RILs homozygous for the cv. Paragon *NAM-A1* (non-staygreen) allele, *P=*0.002, whilst no differences relating to *NAM-A1* were observed, *P=*0.55. Furthermore, mean grain width and TGW of ‘early’ heading *Ppd* x 1189a RILs homozygous for the *NAM-A1* variant were lower compared to those homozygous for the cv. Paragon allele in Germany in 2018, *P=*0.02, and in the UK in 2016, albeit not significantly, *P≤*0.2 (Fig. 8). Grain samples were unavailable for trials conducted in France, preventing grain morphometric assessment.

Combined, multi-environmental testing of *Ppd* x staygreen RILs does not support our hypothesised synergistic, grain yield enhancing, relationship between early heading and staygreen traits (Fig. 1). Instead, phenological, not source related, traits remained a greater determinant of final grain yield. Indeed, grain morphometric analysis reveals that combining *Ppd-1a* and *NAM-A1* allelic variants can reduce grain size, suggesting trait antagonism.

### Combining *Ppd-1a* and *NAM-A1* variants increases competition for resource delivery to the grain

For Paragon x staygreen populations and ‘late’ heading *Ppd* x staygreen RILs, *NAM-1* variants were associated with increasing final grain yield via enhancing mean grain width and TGW (Fig. 3). Analysis of grain yield components confirmed our *NAM-1* variants influenced grain size, not grain number, however TGW of ‘early’ *Ppd* x staygreen RILs is typically even greater (Fig. 8). While grain fill duration of 1189a and 2316b mutants is longer than parental cv. Paragon (Chapman *et al*., 2021b), so is grain fill duration of *Ppd-1a* versus *Ppd-1b* carrying lines (Novoselović *et al*., 2015; Royo *et al*., 2018). Yet, the introduction of *NAM-1* variation into a *Ppd-1a* background does influence grain filling dynamics, with variation in grain maturity observed across all locations amongst ‘early’ heading *Ppd* x staygreen RILs contrasting for *NAM-1* alleles. When visiting Germany on 22^nd^ June 2018, despite plants being close to terminal senescence grains of *Ppd* x 1189a RILs homozygous for the *NAM-A1* variant remained soft, whilst those carrying the cv. Paragon allele were mature. Such asynchronous development suggests the benefits associated with staygreen phenotypes, including greater photosynthate availability and grain filling extension, are unrealised in *Ppd-1a* conferred ‘early’ heading backgrounds. Moreover, ‘early’ heading and staygreen traits may prove antagonistic, with resources destined for the grain retained, promoting plant survival and delaying senescence.

To investigate the physiological effect of combining *Ppd-1a* and *NAM-1* variation, grain filling experiments were conducted for ‘early’ heading *Ppd* x staygreen RILs contrasting for *NAM-1* variants in summer 2019. RILs selected were *Ppd* x 1189a-56 and *Ppd* x 2316b-57, alongside triple *Ppd-1a* parental NILs for comparison, and were part of the subsets included in multi-environment experiments (Supplementary Table S1). To assess grain filling dynamics, grain moisture and grain weight were recorded from anthesis to maturation, alongside flag leaf and peduncle senescence to confirm senescence phenotype (Fig. 9). Grain filling experiments were also performed for parental lines 1189a, 2316b and cv. Paragon, to validate previous results (Chapman *et al*., 2021b; Supplementary Fig S3).

Visualising senescence progression demonstrates the greater penetrance of the 1189a (*NAM-A1*) compared to 2316b (*NAM-D1*) phenotype in an ‘early’ heading background (Fig. 9; Table 5). Flag leaf and peduncle senescence of *Ppd* x 1189a-56 was delayed compared to its respective parental triple *Ppd-1a* NIL, with differences in peduncle phenotype significant, *P*=0.03. Inconsistencies in flag leaf and peduncle senescence were observed for *Ppd* x 2316b-57, suggestive of a ‘non-staygreen’ (leaf) and ‘staygreen’ (peduncle) phenotype (Fig. 9), explained by pre-emergence frost and foliar damage. Once initiated, the senescence rate of *Ppd* x 2316b-57 is slower relative to its respective triple *Ppd-1a* parent (Fig. 9). However, in 2019 the 2316b phenotype was also less penetrative compared to other years (Chapman *et al*., 2021b), not significantly differing from parental cv. Paragon, *P≥*0.19 (Table 5).

Differences in grain filling dynamics between *Ppd* x 1189a-56 and *Ppd* x 2316b-57 and their respective parental triple *Ppd-1a* NILs were typically significant for grain moisture content, *P≤*0.06 (Table 5; Fig. 9). Differences in grain moisture dynamics between *Ppd* x 1189a-56 and parental triple *Ppd-1a* NIL were like those for 1189a and cv. Paragon, with significant differences during the latter stages of grain filling (Fig. 9, Supplementary Table S5). On 2^nd^ July 2019, ∼42 daa, grain moisture content of *Ppd* x 1189a-56 was 9% greater compared to the parental triple *Ppd-1a* NIL, *P=*0.01. On 15^th^ July 2019, this difference increased to +14.5%, *P<*0.0001, with grain moisture content (%) measuring 41.0 [39.2, 42.8] (mean [CI_95%_]); approximately the time that rapid grain filling ceases (Neghliz *et al*., 2016). Conversely, between 10^th^ and 19^th^ July 2019, ∼50-59 daa, grain moisture content of *Ppd* x 2316b-57 was significantly lower compared to the parental triple *Ppd-1a* NIL, *P≤*0.002 (Fig. 9). These results support rejection of our original hypothesis (Fig. 1), as combining ‘early’ heading with staygreen traits fails to further extend grain fill duration, at least for 2316b (*NAM-D1*).

Regarding grain weight accumulation, differences were observed during the early-mid grain filling period (Fig. 9). On 19^th^ June 2019, 35 daa, dry grain weight of *Ppd* x 1189a-56 was 27.1% greater relative to the parental triple *Ppd-1a* NIL, *P=*0.01 (Fig. 9; Supplementary Table S5). Similarly, between 19^th^ and 24^th^ June (∼35 to 40 daa) dry grain weight of *Ppd* x 2316b-57 was between 16.9 to 25.2% greater relative to its parental triple *Ppd-1a* NIL, *P≤*0.02. However, towards the end of grain filling between 19^th^ and 25^th^ July, dry grain weight of *Ppd* x 2316b-57 was between 13.7 and 22.8% lower compared to the parental triple *Ppd-1a* NIL, *P≤*0.0007 (Fig. 9, Supplementary Table S5). Differences in dry grain weight between *Ppd* x 1189a-56 and the respective parental triple *Ppd-1a* NIL were only significant on 25^th^ July, *P=*0.03 (Fig. 9; Supplementary Table S5). Due to the low grain moisture content, dry grain weights recorded on July 25^th^, 2019, provide an estimate of TGW, with values significantly lower for *Ppd* x 1186-56 and *Ppd* x 2316b-57 compared to parental triple *Ppd-1a* NILs, *P*≤0.03 (Fig. 9, Supplementary Table 5). Thus, the earlier staygreen trait-associated increases in grain weight went unrealised regarding final grain weight. Results obtained for *Ppd* x 1189a-56 concur with those for Germany, whereupon the *NAM-A1* mutation is associated with significantly reducing TGW in an ‘early’ heading background, *P=*0.02 (Fig. 8). Combined, this suggests the additional photosynthates associated with the 1189a *NAM-A1* variant are retained in the straw, with the cv. Paragon *NAM-A1* allele associated with greater remobilisation efficiency.

### Variation in grain protein content relates to extremity of staygreen phenotype

Grain yield and protein content are negatively correlated (Simmonds, 1995). *NAM-1* is a positive regulator of senescence, grain protein, and mineral content (Uauy *et al*. 2006a; 2006b). Our *NAM-1* variants are associated with a reduction in grain protein content proportional to senescence phenotype severity (Supplementary Table S6; Supplementary Fig. S6). When *NAM-1* variants are associated with increasing final grain yield (2017, *P*≤0.06; Fig. 3), differences emerge concerning grain protein deviation. For Paragon x 1189a RILs, the correlation between grain protein content and grain yield was r=-0.89, *P*<0.0001, and r=-0.7, *P*<0.003, for those homozygous for the *NAM-A1* mutant and cv. Paragon allele respectively. For Paragon x 2316b RILs, correlations of r=-0.39, *P*=0.053, and r=-0.63, *P*=0.007, were reported for RILs homozygous for the *NAM-D1* variant and cv. Paragon allele respectively (Supplementary Fig. S6). The less negative correlation associated with the 2316b (*NAM-D1*) staygreen trait is indicative of positive grain protein deviation, which results of Mosleth *et al*. (2020) corroborate. In 2018, correlations between grain protein content and grain yield for Paragon x staygreen RILs were typically not significant, *P*=0.02 to 0.8 (Supplementary Fig. S6).

## Discussion

### Confirming Utility of Staygreen Traits Under Maritime Conditions

Contradictory relationships between staygreen traits and yield exist (Gregersen *et al*., 2013). Kipp *et al*. (2014) report a negative relationship between onset of senescence and final grain yield, *r^2^*=0.81, *P*≤0.01, whilst under water stress Moraga *et al*., 2022 report a positive correlation, *R*=0.54-0.62, *P*<0.01. For a winter wheat diversity set comprising >300 cultivars grown in Switzerland, Anderegg *et al*. (2020) report Pearson correlations between onset of senescence and grain yield of *r*=0.173-0.369, *P*<0.01. Delayed senescence is often associated with stress adaptation (Vijayalakshmi *et al*., 2010; Gregersen *et al*., 2013; Thomas and Ougham, 2014; Jagadish *et al*., 2015), which our results confirm. Under moderate heat stress and water limitation we report a staygreen trait associated grain yield increase of up to 8.5% (Fig. 3), identifying no yield penalty in the absence of stress (Fig. 3; Fig. 5; Fig. 7).

Our staygreen trait-associated grain yield increases result from greater mean grain width and TGW, proportional to the delays in senescence, and grain filling extension, observed for 1189a (*NAM-A1*) and 2316b (*NAM-D1*) mutants (Chapman *et al*., 2021b) and derived populations (Chapman *et al*., 2021a). Globally, the UK wheat growing season is one of the longest due to the oceanic climate and relatively low ambient temperatures (Semenov *et al*., 2014; Mueller *et al*., 2015), not renowned for late-season heat and drought stress under which staygreen traits confer tolerance, making staygreen trait-associated increases in final grain yield potentially unexpected (Fischer, 2011). Results for Paragon x 1189a RILs accord with Alhabbar *et al*. (2018). Here, the *NAM-A1* variant encoded by cv. Mace was associated with an 8-day grain filling extension and 13-day delay in maturity relative to cv. Westonia (Alhabbar, Islam, *et al*., 2018). A study concerning *NAM-A1* and *NAM-B1* allelic combinations also report a positive correlation between their *NAM-A1* variant and TGW, *r*=0.31, *P*<0.05, equating to a 1.87% increase (Alhabbar, Yang, *et al*., 2018), lower than reported here (Fig. 3). However, our results contradict Cormier *et al*. (2015) who report a *NAM-A1* associated TGW reduction, *P<*0.05, alongside Avni *et al*. (2014) and Leonova *et al*. (2022), who found no difference, *P*>0.05. We conclude combinations of these *NAM-A1* and *NAM-B1* missense variants deliver high yield potential associated with grain fill extension, similar to Alhabbar *et al*. (2018). Results for the 2316b staygreen trait enable extension of this hypothesis, whereupon our NAM-D1 variant is associated with grain filling extension (Chapman et al., 2021b) and TGW improvement (Fig. 3; Fig. 7).

In the UK in 2017 and 2018, staygreen trait-associated grain width increases improved grain yield. The elevated temperatures of 2018 significantly impact grain filling, with every 1 °C above 15-20 °C equating to a 1.5 mg day^-1^ reduction in grain weight (Dias and Lidon, 2009; Farooq *et al*., 2011). Such stress resulted in source-limitation, with senescence initiated earlier, progressing rapidly, and grain fill curtailment, which our *NAM-1* variants countered. Summer 2016 was notable for its absence of stress, characterized by mild temperatures and surplus rainfall (Supplementary Fig. S1), delaying senescence. Data for *Ppd* x staygreen RILs visually demonstrates this, whereupon senescence curves of ‘early’ and ‘late’ heading RILs converge upon one another respective to senescence type (Fig. 4; Supplementary Fig. S2). TGW of ‘early’ & ‘late’ heading *Ppd* x 2316b RILs was similar, *P*=0.52, whilst a heading-senescence interaction was observed for *Ppd* x 1189a RILs, *P*=0.034 (Fig. 8). Therefore, under such conditions the effect of these staygreen traits on grain yield is, at worst, neutral, unless sink size, grain length or number, is increased.

Weather fluctuations observed between 2016 and 2018 are increasingly common under climate change, making adoption of grain fill duration and staygreen traits key to improving drought tolerance (Hammer *et al*., 2010; Farooq *et al*., 2011; Christopher *et al*., 2016; Senapati *et al*., 2019). Using the Sirius wheat model to predict performance of existing cultivars and identify wheat ideotypes optimised for 2050 climate change conditionside, Semenov *et al*. (2014) found grain fill duration and maturation of optimised cultivars as 2-3 weeks longer, and 19 days later, respectively. Our results support *NAM-1* variant adoption as a hedge-betting strategy, contributing to grain yield increases under UK-type stress conditions without introducing significant yield penalties under favourable conditions (Fig. 3).

### Physiological Trade-offs: Source to sink dynamics and the influence of *Ppd-1*

Greater TGW of Paragon x staygreen RILs homozygous for *NAM-1* variants was driven by grain width (Fig. 3). For *Ppd* x staygreen RILs this was unobserved (Fig. 8), with differences in final grain yield due to phenology (Fig. 5; Supplementary Table S4). An absence of synergy between earlier heading and staygreen trait-associated grain filling extension could represent a physiological trade off (Fig. 1), necessitating complementary source and sink trait manipulation (Abbai *et al*., 2023; Murchie *et al*. 2023).

Final grain yield represents a trade-off between grain number and grain size. The significantly lower spikelet (& grain) numbers of ‘early’ compared to ‘late’ heading *Ppd* x staygreen RILs were compensated by their greater TGW (Fig. 8; Supplementary Table S4). Between 2016 and 2018, TGW of ‘early’ heading *Ppd* x staygreen RILs was 16.5 ±12.6 % (mean ± SD) greater than their ‘late’ heading counterparts (Fig. 8). In 2018, under increasing water limitation and heat stress (Supplementary Fig. S1), TGW of ‘early’ heading *Ppd* x 1189a RILs approached the theoretical maximum of ∼61.4 g (Rasheed *et al*., 2014).

In the UK in 2017 and 2018, *NAM-1* variants were associated with increasing final grain yield of Paragon x staygreen RILs, suggesting source-limitation. These results build on our previous work, whereby delays in senescence onset of 1189a and 2316b led to proportionate grain fill extensions (Chapman *et al*., 2021b). Grain maturity assessment of associated segregating RIL populations confirmed this (Chapman *et al*., 2021a), demonstrating these observed differences translated into grain yield improvement.

For *Ppd* x staygreen RILs, staygreen trait-associated differences in grain maturation were observed in France & Germany, but not grain yield (Fig. 5). Moreover, grain size of ‘early’ heading *Ppd* x 1189a RILs homozygous for the *NAM-A1* allele was lower in the UK in 2016 and Germany in 2018, indicating a reduction in leaf to grain resource remobilization. This indicates competition for resources between live tissues and the grain, or differences in starch deposition, which Jenner *et al*. (1991) attributes grain weight variation to. Here, the known *NAM-1* variant associated reduction in remobilisation efficiency (Avni et al., 2014) was likely compounded by *Ppd-1a* (Royo *et al*., 2018).

Avni et al. (2014) report flag leaf and peduncle nitrogen content of a combined *gpc-a1* and *gpc-d1* mutant as 73% and 57% greater (*P*<0.001, 50 DAA), respectively, with grain nitrogen 37 % lower than the control (*P*<0.01, 60 DAA). Whilst, a study of 34 spring durum wheat cultivars contrasting for *Ppd-1* composition reports translocation efficiency of pre-flowering assimilates as 24% greater for those carrying *Ppd-1b* compared to *Ppd-1a* alleles (Royo *et al*., 2018).

### How to reap the benefits associated with staygreen traits in earlier flowering backgrounds

Through *Ppd-1a* allelic introgression, we developed RILs contrasting in *NAM-1* allelic composition, engineered to assess heading-date independent staygreen trait expression (Fig. 4). Combining *Ppd-1a* and *NAM-1* mutant alleles did produce earlier flowering staygreen types but increased sink-limitation. The *Ppd-1a* associated pleiotropic reduction in grain number, combined with the inherently longer grain filling period, reduced penetrance of the staygreeen trait-associated grain filling extension.

Regarding the 1189a (*NAM-A1*) phenotype, *Ppd-1a* allele adoption disrupted source:sink dynamics, explaining the ‘ripe ear, green plant’ phenotype observed in 2016 (Chapman *et al*., 2021a). Relative to 2316b, the severity of the delay in senescence onset for 1189a represents greater inter-organ resource competition, explaining the differential trends in TGW and mean grain width reported for respective *Ppd* x staygreen RILs.

Given the negative pleiotropic interaction observed between *Ppd-1a* and *NAM-1* variants for *Ppd* x staygreen RILs, unlocking the potential benefits associated with *NAM-1* variants may require more sophisticated phenological manipulation. One strategy would be combining *Ppd-D1a* and *TaTOE-B1* (Avalon-type) alleles, as explored by Dreisigacker *et al*. (2021) when reporting effects of flowering related genes on yield components for two high biomass spring wheat panels. Here, *Ppd-D1a* had the most significant positive effect on yield via increasing biomass at plant maturity and harvest index. Conversely, *TaTOE-B1* influences grain number, with the Avalon type allele associated with a 4.7-5% greater yield compared to the Cadenza type, due to increases in fruiting efficiency, spikelets spike^-1^, and grain filling rate (Dreisigacker *et al*., 2021).

### Remobilising, and realising potential of, staygreen-trait associated additional photosynthates

Between 2016 and 2018 both source and sink limitation were observed. Previously, Borrill *et al*. (2015) reported the additional photosynthates produced by *NAM* RNAi lines were stored as stem fructans, and not remobilized. To counteract this, *NAM-1* variants could be coupled with the cv. Westonia *1-FEH-w3* allele (Zhang *et al*., 2015). *1-FEH-w3* is a major regulator of stem fructans remobilization, with the promoter deletion encoded by cv. Westonia associated with increased gene expression, TGW and grain number per spike under drought, P<0.05 (Zhang *et al*., 2015).

Alternative strategies could involved joint targeting of staygreen and remobilisation traits, including identification of differentially expressed genes within peduncle tissue (Taria *et al*., 2025), or further manipulation of source-sink strength, via improved spike branching to increase grain number (Abbai *et al*., 2023). Moreover, targetted engineering of Rubisco proteins could alter carbon-nitrogen dynamics (Paul *et al*., 2026), which could counter the differential gene expression associated with *NAM-1* under contrasting nitrogen regimes (Andleeb *et al*., 2023).

Thus, combining staygreen and remobilisation traits could (re-)balance source:sink demands, enabling realization of potential staygreen-trait associated benefits under wide ranging environments.

## Conclusion

Combining *Ppd-1* and *NAM-1* variants enabled successful dissection, and uncoupling, of heading & senescence trait relationships. Regarding grain yield, *Ppd-1a* allelic adoption resulted in earlier flowering but reduced sink size (grain number), which the staygreen-trait associated increase in source, and grain size, could not compensate. In ‘late’ heading (*Ppd-1b*) backgrounds, we demonstrated the potential utility of our *NAM-1* variants. Under moderate stress, staygreen traits increased final grain yield, with the observed extension in grain filling duration becoming realized as significant increases in mean grain width, improving TGW. Here, we report the agronomic utility of a *NAM-D1* variant for the first time. *NAM-D1* associated grain yield improvements were typically lower compared to *NAM-A1*, but not at the cost of agronomic acceptability. Combined, staygreen trait adoption requires more fine-tuned and targeted approaches to realise full trait potential.

## Supplementary Material

**Supplementary Fig. S1.** Daily rainfall and mean daily temperature data

**Supplementary Fig. S2.** Senescence of Paragon x staygreen RILs plotted against thermal time

**Supplementary Fig. S3.** Senescence of *Ppd* x staygreen RILs (Norwich, 2016-2018)

**Supplementary Fig. S4.** Senescence of *Ppd* x staygreen RILs plotted against thermal time

**Supplementary Fig. S5.** Results of grain filling experiments for 1189a, 2316b & cv. Paragon (2019)

**Supplementary Fig. S6.** Grain protein content deviation of Paragon x staygreen RILs (Norwich, 2017-2018)

**Supplementary Table S1.** *Ppd* x staygreen RIL selections & trial overview

**Supplementary Table S2.** Yield components for Paragon x staygreen trials (2016-2018)

**Supplementary Table S3.** Tukey *post-hoc* test results concerning senescence of *Ppd* x staygreen RIL subsets (UK, 2016-2018).

**Supplementary Table S4.** Yield components for *Ppd* x staygreen trials (2016-2018)

**Supplementary Table S5.** Differences in grain filling parameters, pairwise-comparison (2019)

**Supplementary Table S6.** Grain protein content of Paragon x staygreen & *Ppd* x staygreen RILs (2016-2018)

## Acknowledgements

The authors wish to thank the JIC field experimentation team, without whom field trials would not have been possible, alongside horticultural services and glasshouse staff. Thanks go to field trial teams at KWS-UK, Thriplow, KWS Momont, France, and KWS Lochow, Germany, who made multi-site experiments a reality, collected invaluable data, and assisted Elizabeth when visiting sites. Special thanks go to Nick Bird, Anne-Marie Brosnan, Emile, Thea and Yeorgia for logistics and practical assistance. Thanks go to Luzie Wingen, James Brown, and Richard Morris for statistical support, and Rajani Awal and Richard Goram for tissue collection and DNA extraction. Additional thanks go to Gemma Molero, whereby our discussions helped interpretation of grain filling results and associated physiological concepts. The original mutants were developed by Robert Koebner and Leodie Alibert. E.A.C. dedicates this publication in memory of Philippa Borrill, whose death represents a significant loss to the wheat community, and an invaluable intellectual sparring partner.

## Authors’ Contributions

E.A.C., J.L., S.G. conceived the study. S.O. screened, developed, and maintained the genetic material. E.A.C. and J.L. organised, designed, and conducted field experiments. R.B. performed grain filling experiments conducted in 2019 under the supervision of E.A.C. S.G., J.L., and S.O. provided supervision and technical assistance for E.A.C. E.A.C. analysed the data and wrote the article in correspondence with all authors.

## Data Availability

Germplasm used in the current study have been deposited with the Germplasm Resources Unit (GRU) at John Innes Centre. The datasets generated during and/or analysed during the current study are available from the corresponding author upon reasonable request.

