## Supplementary Material for "Extending the seasons at both ends? Understanding the physiological and genetic context required for stay green mediated yield increase in wheat (*Triticum aestivum*)"

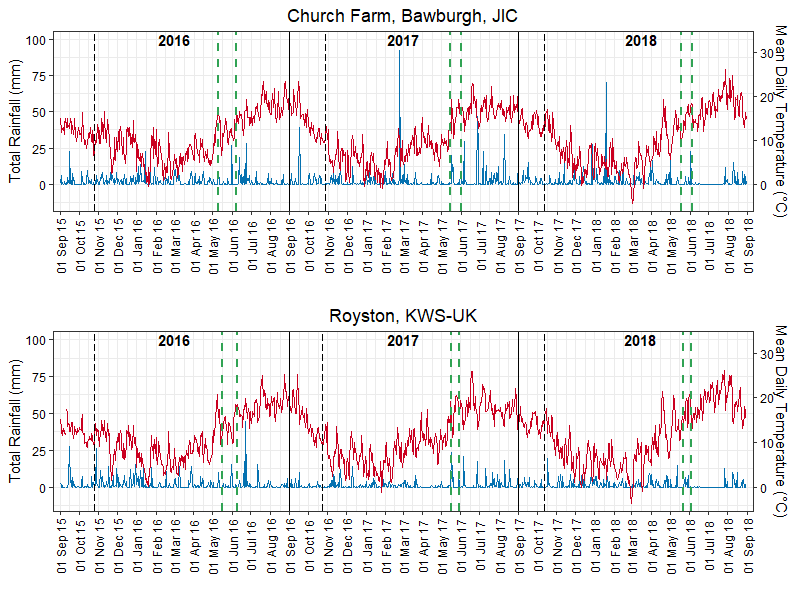

**A**

**B**

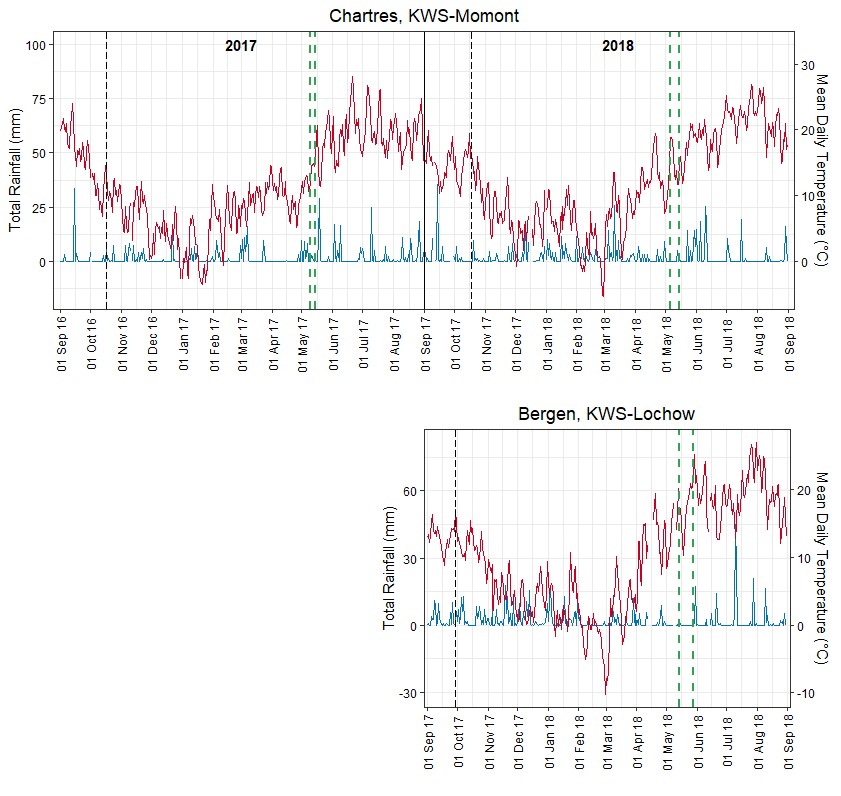

**C**

**D**

**Figure S1: Daily rainfall (mm) and mean daily temperature (°C) for trial sites between 2016 to 2018.** **(A)** Cburch Farm, Bawburgh, Norwich (JIC), **(B)** Cambridgeshire, KWS-UK, **(C)** Chartes, France, KWS-Momont, **(D)** Bergen, Germany KWS-Lochow. Y-axes refer to daily total rainfall (mm) (red), and mean daily temperature (°C) (blue), left and right respectively. Black dotted lines indicate drilling date. Green dashed lines indicate ear emergence (GS55) window (From *Ppd-1a* to standard cultivar for the region).

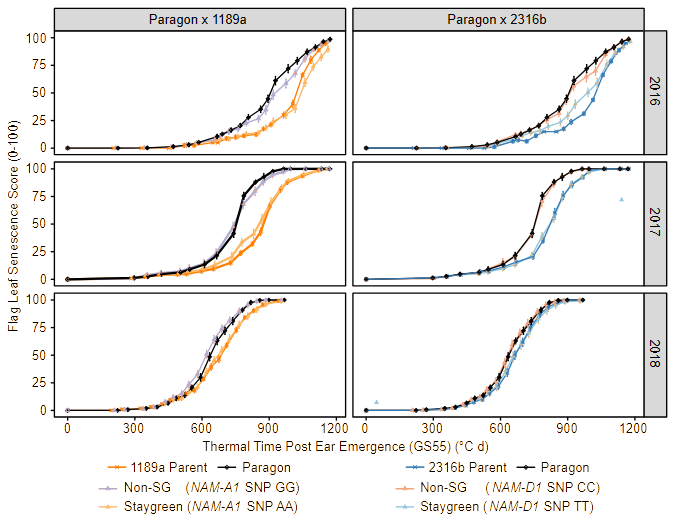
**Figure S2: Senescence progression of Paragon x mutant RILs when plotted against thermal time (°C day).** Progression of flag leaf senescence of Paragon x 1189a (right) and Paragon x 2316b (left) RILs when grouped by *NAM-1* composition, plus parental lines (Paragon, black; 1189a, dark orange; 2316b, dark blue), Norwich, 2016-2018. ‘Non-staygreen’ (Non-SG; purple, 1189a; red, 2316b) and ‘staygreen’ (orange, 1189a: blue, 2316b) refers to whether RILs are homozygous for the mutant or wildtype allele, respectively. Senescence was scored visually using a 0-100 scale 3-4 times per week from ear emergence (GS55), and scoring date converted to thermal time (°C day). Mean±SEM, n≥15 per allelic group, n=1-3 per RIL.

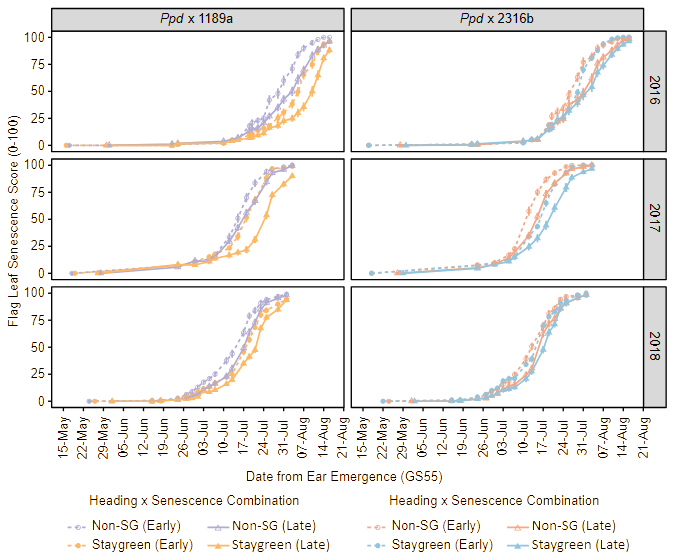

**Figure S3: Staygreen trait expression for ‘early’ and ‘late’ heading *Ppd* x staygreen RILs, Norwich 2016-2018.** Heading type, ‘early’ (circles, dotted lines), ‘late’ (triangles, solid lines). Senescence type, ‘non-staygreen’ (Non-SG; purple, 1189a; red, 2316b), ‘staygreen’ (orange, 1189a; blue, 2316b). ‘Staygreen’ refers to homozygosity for either *NAM-1* variant (*NAM-A1,* 1189a, or *NAM-D1,* 2316b), with the ‘non-staygreen’ group homozygous for the cv. Paragon allele. Senescence was scored visually using a 0-100 scale three to four times per week from ear emergence (GS55), mean ± SEM, n≤15 per heading-senescence combination, n=1-3 per RIL.

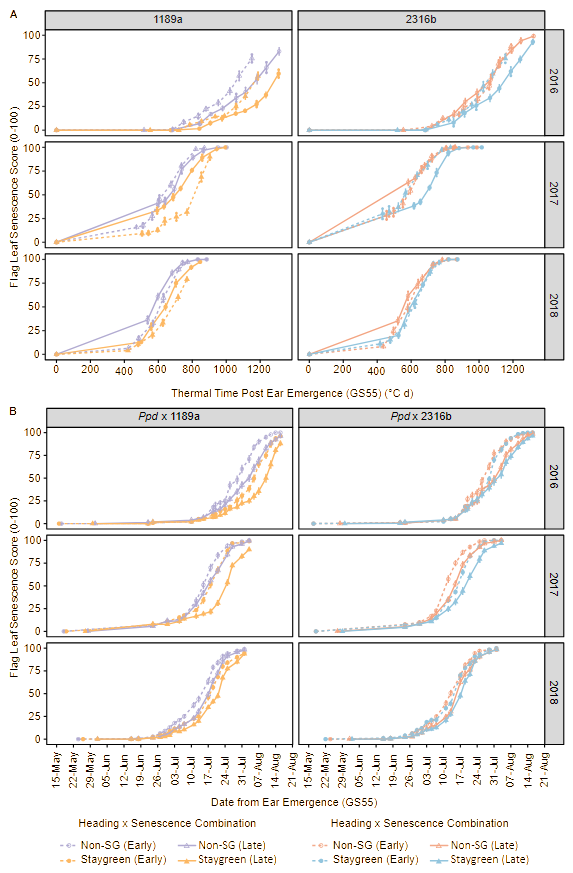

**Figure S4:** **Staygreen trait expression for ‘early’ and ‘late’ heading *Ppd* x staygreen RILs adjusted for heading date variation.** Flag leaf senescence progression of *Ppd* x staygreen RILs plotted against thermal time when grown in Cambridgeshire (A), and Norwich (B), 2016-2018. Heading type, ‘early’ (circles, dotted lines), ‘late’ (triangles, solid lines). Senescence type, ‘non-staygreen’ (Non-SG; purple, 1189a; red, 2316b), ‘staygreen’ (orange, 1189a; blue, 2316b). ‘Staygreen’ refers to homozygosity for either *NAM-1* variant (*NAM-A1,* 1189a, or *NAM-D1,* 2316b), with the ‘non-staygreen’ group homozygous for the cv. Paragon allele. Senescence was scored visually using a 0-100 scale two to four times per week from ear emergence (GS55), with scoring dates converted into thermal time (day °C). Mean ± SEM, n≤15 per heading-senescence combination, n=1-3 per RIL.

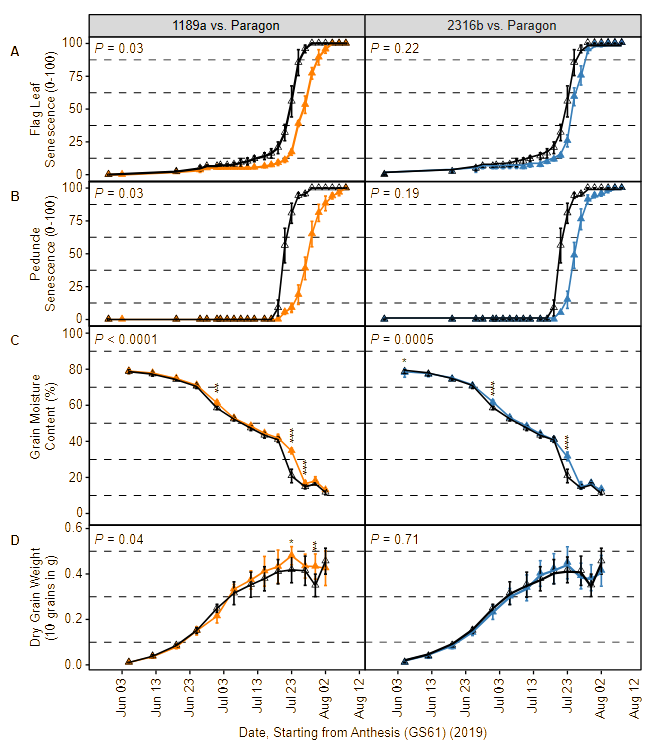
**Figure S5: Confirming delayed senescence is associated with grain filling extension.** Grain filling experiments conducted for 1189a (*NAM-A1*; left, orange) and 2316b (*NAM-D1*, right, light blue), alongside parental cv. Paragon (black), Norwich, 2019. (A) Visual leaf senescence and (B) Peduncle senescence; 0-100 scale, mean ± SD. Senescence was scored at the plot level every 2-4 days following anthesis (GS61), n=3. (C) Grain moisture content (%) and (D) Dry grain weight (10 grains in g), were recorded to determine grain filling dynamics, mean ± SD. 4-5 ears were sampled per plot every 4-6 days from anthesis, 2 plots per genotype. *P*-values represent overall differences (Table 2). Differences at specific time points according to Tukey *post-hoc* tests are indicated. *P-*values, *<0.05, **<0.01, ***<0.0001.

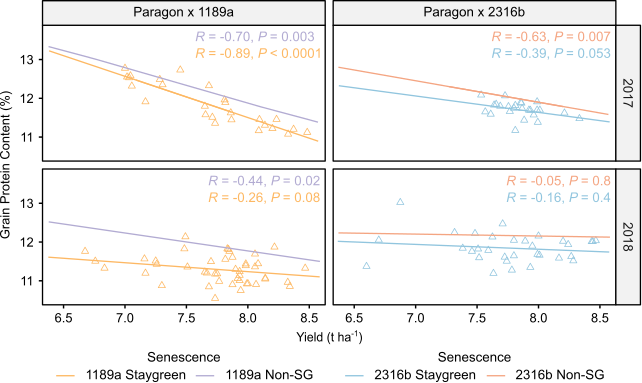

**Figure S6:** **Senescence can influence the relationship between grain protein content & final grain yield**. Grain protein content (%) compared to final grain yield (t ha^-1^) of Paragon x 1189a (right) and Paragon x 2316b (left) RILs, Norwich, 2017 (top) & 2018 (bottom). Significant differences between years. Spearman’s rank correlations calculated using mean values of each RIL, n=3 (2017), n=2 (2018). No. of RILs ≥ 42 per population. Staygreen RILs (homozygous for *NAM-1* variants), orange (1189a, *NAM-A1*), blue (2316b, *NAM-D1*). Non-staygreen RILs (homozygous for cv. Paragon allele), purple (1189a), red (2316b).

**Supplementary Tables**

**Table S1: Summary of *Ppd* x staygreen RILs trialled between 2016-2019.**

| ***Ppd*  x 1189a RIL** | **Year** | | | | **2016** | | | | **2017** | | | | | | **2018** | | | | | | | |
| --- | --- | --- | --- | --- | --- | --- | --- | --- | --- | --- | --- | --- | --- | --- | --- | --- | --- | --- | --- | --- | --- | --- |
|  | **Location, Plot size & replication** | | | | **JIC** | | **KWS-UK** | | **JIC** | | **KWS-UK** | | **France** | | **JIC** | | **KWS-UK** | | **France** | | **Germany** | |
|  | **Heading Class** | | ***NAM-A1* genotype*** | | 1x1 m | | 1x6 m | | 2x1 m | | 2x6 m | | 3x7.5 m | | 2x1 m | | 3x7 m | | 2-3x7.05 m | | 3x5 m | |
| Ppdx1189a-4 | Late | | A | | x | | x | |  | |  | |  | |  | |  | |  | |  | |
| Ppdx1189a-5 | Late | | H | | x | | x | | x | | x | |  | | x | | x | |  | |  | |
| Ppdx1189a-11 | Late | | A | | x | | x | | x | | x | |  | | x | | x | |  | |  | |
| Ppdx1189a-13 | Early | | B | | x | | x | |  | |  | |  | |  | |  | |  | |  | |
| Ppdx1189a-15 | Late | | B | | x | | x | |  | |  | |  | |  | |  | |  | |  | |
| Ppdx1189a-16 | Early | | A | | x | | x | | x | | x | | x | | x | | x | | x | | x | |
| Ppdx1189a-18 | Late | | A | | x | | x | | x | | x | |  | | x | | x | |  | |  | |
| Ppdx1189a-20 | Early | | A | | x | | x | | x | | x | | x | | x | | x | | x | | x | |
| Ppdx1189a-22 | Early | | A | | x | | x | | x | | x | | x | | x | |  | |  | |  | |
| Ppdx1189a-26 | Early | | B | | x | | x | |  | |  | |  | |  | |  | |  | |  | |
| Ppdx1189a-27 | Late | | A | | x | | x | |  | |  | |  | |  | |  | |  | |  | |
| Ppdx1189a-31 | Late | | A | | x | | x | | x | | x | |  | | x | |  | |  | |  | |
| Ppdx1189a-32 | Late | | B | | x | | x | |  | |  | |  | |  | |  | |  | |  | |
| Ppdx1189a-35 | Late | | A | | x | | x | | x | | x | |  | | x | | x | |  | |  | |
| Ppdx1189a-36 | Early | | - | | x | | x | |  | |  | |  | |  | |  | |  | |  | |
| Ppdx1189a-40 | Early | | B | | x | | x | | x | | x | | x | | x | |  | |  | |  | |
| Ppdx1189a-42 | Late | | A | | x | | x | | x | | x | |  | | x | |  | |  | |  | |
| Ppdx1189a-46 | Late | | B | | x | | x | |  | |  | |  | |  | |  | |  | |  | |
| Ppdx1189a-47 | Early | | B | | x | | x | | x | | x | | x | | x | | x | | x | | x | |
| Ppdx1189a-48 | Late | | B | | x | | x | | x | | x | |  | | x | | x | |  | |  | |
| Ppdx1189a-52 | Late | | B | | x | | x | | x | | x | |  | | x | | x | |  | |  | |
| Ppdx1189a-56 | Early | | B | | x | | x | | x | | x | | x | | x | | x | | x | | x | |
| Ppdx1189a-58 | Early | | B | | x | | x | |  | |  | |  | |  | |  | |  | |  | |
| Ppdx1189a-59 | Early | | B | | x | | x | |  | |  | |  | |  | |  | |  | |  | |
| Ppdx1189a-60 | Early | | B | | x | | x | |  | |  | |  | |  | |  | |  | |  | |
| Ppdx1189a-61 | Early | | A | | x | | x | | x | | x | | x | | x | | x | | x | | x | |
| Ppdx1189a-62 | Early | | B | | x | | x | |  | |  | |  | |  | |  | |  | |  | |
| Ppdx1189a-64 | Late | | B | | x | | x | |  | |  | |  | |  | |  | |  | |  | |
| Ppdx1189a-65 | Early | | B | | x | | x | |  | |  | |  | |  | |  | |  | |  | |
| Ppdx1189a-70 | Late | | B | | x | |  | |  | |  | |  | |  | |  | |  | |  | |
| Ppdx1189a-72 | Late | | A | | x | | x | |  | |  | |  | |  | |  | |  | |  | |
| Ppdx1189a-75 | Early | | B | | x | | x | |  | |  | |  | |  | |  | |  | |  | |
| Ppdx1189a-79 | Late | | A | |  | | x | |  | |  | |  | |  | |  | |  | |  | |
| Ppdx1189a-80 | Late | | B | | x | | x | |  | |  | |  | |  | |  | |  | |  | |
| Ppdx1189a-81 | Early | | B | | x | | x | |  | |  | |  | |  | |  | |  | |  | |
| Ppdx1189a-83 | Late | | H | | x | | x | |  | |  | |  | |  | |  | |  | |  | |
| Ppdx1189a-92 | Early | | A | | x | | x | | x | | x | | x | | x | | x | | x | | x | |
| ***Ppd*  x 2316b RIL** | | **Year** | | | | **2016** | | | | **2017** | | | | | | **2018** | | | | | | |
|  |  | **Location, Plot size & replication** | | | | **JIC** | | **KWS-UK** | | **JIC** | | **KWS-UK** | | **France** | | **JIC** | | **KWS-UK** | | **France** | | **Germany** |
|  |  | **Heading Class** | | ***NAM-D1* genotype*** | | 1x1 m | | 1x6 m | | 2x1 m | | 2x6 m | | 3x7.5 m | | 2x1 m | | 3x7 m | | 2-3x7.05 m | | 3x5 m |
| Ppdx2316b-1 | | Early | | B | | x | | x | |  | |  | |  | |  | |  | |  | |  |
| Ppdx2316b-2 | | Early | | H | | x | | x | |  | |  | |  | |  | |  | |  | |  |
| Ppdx2316b-3 | | Early | | A | | x | | x | | x | | x | | x | | x | |  | |  | |  |
| Ppdx2316b-4 | | Early | | B | | x | | x | |  | |  | |  | |  | |  | |  | |  |
| Ppdx2316b-5 | | Early | | B | | x | | x | |  | |  | |  | |  | |  | |  | |  |
| Ppdx2316b-8 | | Late | | B | | x | | x | | x | | x | |  | | x | | x | |  | |  |
| Ppdx2316b-12 | | Early | | A | | x | | x | | x | | x | | x | | x | | x | | x | | x |
| Ppdx2316b-13 | | Early | | A | | x | | x | | x | | x | | x | | x | | x | | x | | x |
| Ppdx2316b-14 | | Early | | A | | x | | x | | x | | x | | x | | x | | x | | x | | x |
| Ppdx2316b-15 | | Early | | B | | x | | x | |  | |  | |  | |  | |  | |  | |  |
| Ppdx2316b-17 | | Late | | B | | x | | x | | x | | x | |  | | x | |  | |  | |  |
| Ppdx2316b-18 | | Early | | B | | x | | x | | x | | x | | x | | x | | x | | x | | x |
| Ppdx2316b-20 | | Late | | B | | x | | x | | x | | x | |  | | x | | x | |  | |  |
| Ppdx2316b-22 | | Late | | B | | x | | x | |  | |  | |  | |  | |  | |  | |  |
| Ppdx2316b-26 | | Late | | A | | x | | x | | x | | x | |  | | x | | x | |  | |  |
| Ppdx2316b-28 | | Late | | A | | x | | x | | x | | x | |  | | x | | x | |  | |  |
| Ppdx2316b-29 | | Late | | B | | x | | x | |  | |  | |  | |  | |  | |  | |  |
| Ppdx2316b-31 | | Early | | H | | x | | x | | x | | x | | x | | x | | x | | x | | x |
| Ppdx2316b-35 | | Late | | H | | x | | x | | x | | x | |  | | x | |  | |  | |  |
| Ppdx2316b-37 | | Late | | A | | x | | x | |  | |  | |  | |  | |  | |  | |  |
| Ppdx2316b-40 | | Late | | B | | x | | x | |  | |  | |  | |  | |  | |  | |  |
| Ppdx2316b-41 | | Late | | B | | x | | x | |  | |  | |  | |  | |  | |  | |  |
| Ppdx2316b-49 | | Early | | B | | x | | x | |  | |  | |  | |  | |  | |  | |  |
| Ppdx2316b-51 | | Late | | A | | x | | x | |  | |  | |  | |  | |  | |  | |  |
| Ppdx2316b-54 | | Early | | B | | x | | x | |  | |  | |  | |  | |  | |  | |  |
| Ppdx2316b-57 | | Early | | B | | x | | x | | x | | x | | x | | x | | x | | x | | x |
| Ppdx2316b-61 | | Late | | A | | x | | x | |  | |  | |  | |  | |  | |  | |  |
| Ppdx2316b-63 | | Late | | H | | x | | x | |  | |  | |  | |  | |  | |  | |  |
| Ppdx2316b-66 | | Late | | H | | x | | x | |  | |  | |  | |  | |  | |  | |  |
| Ppdx2316b-68 | | Early | | B | | x | | x | | x | | x | | x | | x | |  | |  | |  |
| Ppdx2316b-75 | | Late | | A | | x | | x | | x | | x | |  | | x | | x | |  | |  |
| Ppdx2316b-81 | | Late | | B | | x | | x | |  | |  | |  | |  | |  | |  | |  |
| Ppdx2316b-84 | | Early | | B | | x | | x | |  | |  | |  | |  | |  | |  | |  |
| Ppdx2316b-87 | | Late | | B | | x | | x | | x | | x | |  | | x | | x | |  | |  |
| Ppdx2316b-89 | | Early | | A | | x | | x | |  | |  | |  | |  | |  | |  | |  |

**NAM-1* genotyping: A, homozygous wt; H, heterozygous; B, homozygous mutant; - missing data

**Table S2: Grain morphometrics & yield components recorded for Paragon x staygreen RILs between 2016 & 2018 (Mean [CI_95%_])**

| **Year** | **Paragon x Staygreen Population** | **Senescence** | **Grains m^-2^** | **Mean Grain Area  (mm^2^)** | **Minimum Grain Length (mm)** | **Maximum Grain Length (mm)** | **Mean Grain Length (mm)** | **Minimum Grain Width (mm)** | **Maximum Grain Width (mm)** | **Mean Grain Width  (mm)** |
| --- | --- | --- | --- | --- | --- | --- | --- | --- | --- | --- |
| **2016** | **1189a** | **Non-SG** | - | 19.8  [19.5, 20.1] | 4.45  [3.94, 4.96] | 8.33  [7.78, 8.89] | 6.60  [6.50, 6.64] | 2.21  [1.96, 2.47] | 4.83  [4.55, 5.12] | 3.70  [3.67, 3.72] |
|  |  | **Staygreen** | - | 19.5  [19.2, 19.8] | 4.18  [3.74, 4.62] | 8.03  [7.55, 8.52] | **6.46  [6.40, 6.52]*** | 2.00  [1.78, 2.22] | 4.86  [4.62, 5.11] | 3.69  [3.67, 3.72] |
|  | **2316b** | **Non-SG** | - | 18.9  [18.6, 19.3] | 4.29  [3.79, 4.79] | 8.58  [8.03, 9.13] | 6.47  [6.40, 6.54] | 2.06  [1.82, 2.31] | 4.84  [4.57, 5.12] | 3.59  [3.56, 3.62] |
|  |  | **Staygreen** | - | 18.8  [18.5, 19.0] | 4.29  [3.84, 4.75] | 8.64  [8.14, 9.14] | 6.41  [6.35, 6.48] | 2.19  [1.96, 2.41] | 5.00  [4.75, 5.25] | 3.59  [3.56, 3.61] |
| **2017** | **1189a** | **Non-SG** | 18384  [17946, 18822] | 19.0  [18.7, 19.2] | 4.31  [4.07, 4.55] | 8.45  [8.12, 8.79] | 6.55  [6.48, 6.62] | 2.03  [1.92, 2.14] | 4.92  [4.72, 5.12] | 3.55  [3.52, 3.57] |
|  |  | **Staygreen** | 17898  [17541, 18255] | 19.1  [18.9, 19.3] | 4.04  [3.85, 4.24] | 8.39  [8.13, 8.65] | 6.48  [6.43, 6.54] | 1.99  [1.90, 2.07] | 5.03  [4.87, 5.18] | **3.61  [3.59, 3.63]***** |
|  | **2316b** | **Non-SG** | 20102  [19669, 20535] | 18.2  [17.9, 18.4] | 4.16  [3.93, 4.40] | 8.30  [7.97, 8.62] | 6.42  [6.36, 6.49] | 2.00  [1.90, 2.11] | 4.83  [4.64, 5.02] | 3.47  [3.44, 3.49] |
|  |  | **Staygreen** | 19501  [19100, 19902] | 18.6  [18.4, 18.8] | 4.26  [4.06, 4.45] | 8.36  [8.09, 8.63] | 6.43  [6.37, 6.48] | 1.99  [1.90, 2.08] | 4.91  [4.75, 5.07] | **3.53  [3.51, 3.55]***** |
| **2018** | **1189a** | **Non-SG** | 16090  [15671, 16509] | 17.4  [17.2, 17.6] | 3.32  [3.14, 3.50] | 9.17  [8.84, 9.50] | 6.34  [6.29, 6.38] | 1.53  [1.40, 1.66] | 5.47  [5.25, 5.68] | 3.35  [3.33, 3.36] |
|  |  | **Staygreen** | 16234  [15904, 16564] | **17.9  [17.7, 18.0]***** | 3.41  [3.27, 3.56] | 8.73  [8.47, 8.99] | 6.33  [6.29, 6.36] | 1.67  [1.56, 1.78] | 5.20  [5.02, 5.38] | **3.44  [3.43, 3.46]***** |
|  | **2316b** | **Non-SG** | 17493  [17083, 17903] | 16.7  [16.5, 16.9] | 3.57  [3.41, 3.74] | 9.14  [8.84, 9.45] | 6.25  [6.21, 6.29] | 1.76  [1.64, 1.89] | 5.21  [5.00, 5.41] | 3.25  [3.23, 3.27] |
|  |  | **Staygreen** | 17485  [17149, 17822] | 16.9  [16.8, 17.1] | 3.55  [3.39, 3.71] | 9.09  [8.79, 9.38] | 6.23  [6.19, 6.27] | 1.67  [1.55, 1.79] | 5.33  [5.13, 5.52] | **3.31  [3.29, 3.32]***** |

Staygreen and Non-SG refers to RILs homozygous for *NAM-A1* (1189a) and *NAM-D1* variants, and those homozygous for the cv. Paragon *NAM-1* allele.
Results of pairwise Tukey *post-hoc* comparisons; **P*<0.05; ***P*<0.01, ****P*<0.001 (in bold).

**Table S3: Differences in senescence progression relative to heading group. Results of Tukey *post-hoc* test (*P* value), in accordance with calendar date and thermal time *(italics).***

| **RIL Population** | **Site** | **Senescence** | **2016** | | **2017** | | **2018** | |
| --- | --- | --- | --- | --- | --- | --- | --- | --- |
|  |  |  | **Flag Leaf** | **Peduncle**^†^ | **Flag Leaf*** | **Peduncle** | **Flag Leaf** | **Peduncle** |
| ***Ppd* x 1189a (Early vs. late)** | **Cambridgeshire** | **Non-SG** | 0.031  *(0.56)* | - | <0.0001  *(0.07)* | <0.0001  *(0.006)* | <0.0001  *(<0.0001)* | <0.0001  *(0.0009)* |
|  |  | **Staygreen** | 0.36  *(0.56)* | - | <0.0001  *(<0.0001)* | <0.0001  *(0.006)* | <0.0001  *(<0.0001)* | <0.0001  *(0.0009)* |
|  | **Norwich** | **Non-SG** | 0.06  *(0.0005)* |  | 0.01  *(0.0007)* | 0.0001  *(0.007)* | <0.0001  *(<0.0001)* | 0.0001  *(0.0003)* |
|  |  | **Staygreen** | 0.26  *(0.0005)* |  | <0.0001  *(0.57)* | 0.0001  *(0.87)* | <0.0001  *(<0.0001)* | 0.0001  *(0.0003)* |
| ***Ppd* x 2316b (Early vs. late)** | **Cambridgeshire** | **Non-SG** | <0.0001  *(0.57)* | - | <0.0001  *(0.04)* | <0.0001  *(0.6)* | <0.0001  *(0.0009)* | <0.0001  *(0.05)* |
|  |  | **Staygreen** | 0.0002  *(0.57)* | - | <0.0001  *(0.005)* | 0.0001  *(0.6)* | <0.0001  *(0.0009)* | <0.0001  *(0.05)* |
|  | **Norwich** | **Non-SG** | 0.98  *(<0.0001)* | - | 0.009  *(0.09)* | 0.0004  *(0.21)* | 0.0001  *(<0.0001)* | 0.0003  *(<0.0001)* |
|  |  | **Staygreen** | 0.01  *(<0.0001)* | - | 0.002  *(0.005)* | 0.0004  *(0.004)* | 0.0001  *(<0.0001)* | 0.0003  *(<0.0001)* |

^†^Peduncle senescence was not scored in 2016
*Significant heading-senescence interaction, *P*<0.05

**Table S4: Grain morphometrics & yield components recorded for *Ppd* x staygreen RILs between 2016 & 2018 (Mean [CI_95%_])**

| ***Ppd* x**  **staygreen Population** | **Year** | **Heading** | **Senescence** | **Biomass  0.6 m^-1^ (kg)** | **Harvest Index** | **Spikelets ear^-1^  (mean of 5)** | **Seeds spike^-1^** | **Grains m^-2^** | **Mean Grain area  (mm^2^)** | **Minimum Grain Length (mm)** | **Maximum Grain Length (mm)** | **Mean Grain Length  (mm)** | **Minimum Grain Width (mm)** | **Maximum Grain Width (mm)** | **Mean Grain Width  (mm)** |
| --- | --- | --- | --- | --- | --- | --- | --- | --- | --- | --- | --- | --- | --- | --- | --- |
| **1189a** | **2016** | Early | Non-SG | 0.232  [0.214, 0.249] | **0.40  [0.39, 0.42]^a^** | **19.1  [18.4, 19.7] ^a^** | 54  [51, 57] | 17846  [17044, 18649] | **17.6  [17.1, 18.1] ^a^** | 3.97  [3.26, 4.67] | **8.11  [6.94, 9.29]** | **6.22  [6.12, 6.32] ^a^** | 1.80  [1.49, 2.11] | 5.11  [4.58, 5.63] | 3.41  [3.34, 3.47] |
|  |  |  | Staygreen | 0.254  [0.241, 0.267] | **0.39  [0.38, 0.40]^a^** | **19.4  [18.9, 19.9] ^a^** | 55  [53, 57] | 18002  [17424, 18580] | **17.1  [16.8, 17.3] ^a^** | 3.68  [3.16, 4.19] | **9.98  [9.26, 10.71]*** | **6.23  [6.16, 6.30] ^a^** | 1.66  [1.43, 1.88] | 5.57  [5.20, 5.95] | 3.33  [3.29, 3.37] |
|  |  | Late | Non-SG | 0.238  [0.223, 0.253] | **0.36  [0.35, 0.38^]b^** | **20.8  [20.2, 21.4] ^b^** | 52  [49, 54] | 18867  [18171, 19562] | **15.9  [15.5, 16.2] ^b^** | 3.09  [2.47, 3.71] | 9.84  [8.91, 10.77] | **5.95  [5.86, 6.03] ^b^** | 1.49  [1.22, 1.76] | 5.71  [5.25, 6.17] | 3.27  [3.21, 3.32] |
|  |  |  | Staygreen | 0.261  [0.245, 0.276] | **0.34  [0.33, 0.36]^b^** | **21.2  [20.6, 21.8] ^b^** | 53  [50, 55] | 19022  [18317, 19727] | **16.5  [16.1, 16.8] ^b^** | 2.80  [2.18, 3.42] | 9.48  [8.55, 10.42] | **5.96  [5.88, 6.05] ^b^** | 1.35  [1.08, 1.62] | 6.18  [5.71, 6.64] | 3.32  [3.27, 3.37] |
|  | **2017** | Early | Non-SG | 0.183  [0.168, 0.197] | **0.47  [0.45, 0.50]^a^** | **18.9  [18.1, 19.6] ^a^** | 86  [83, 89] | **14638  [13558, 15718] ^a^** | **17.9  [17.2, 18.7] ^a^** | 4.83  [4.45, 5.20] | 8.26  [7.53, 8.99] | **6.42  [6.30, 6.53] ^a^** | **2.38  [2.19, 2.57] ^a^** | 5.34  [4.89, 5.79] | **3.41  [3.30, 3.51] ^a^** |
|  |  |  | Staygreen | 0.162  [0.145, 0.179] | **0.46  [0.44, 0.49] ^a^** | **19.7  [18.7, 20.6] ^a^** | 88  [84, 92] | **14775  [13474, 16077] ^a^** | **17.8  [16.9, 18.7] ^a^** | 4.61  [4.16, 5.06] | 9.27  [8.39, 10.14] | **6.28  [6.14, 6.42] ^a^** | **2.29  [2.05, 2.53] ^a^** | 5.03  [4.49, 5.57] | **3.46  [3.34, 3.59] ^a^** |
|  |  | Late | Non-SG | 0.179  [0.165, 0.194] | **0.40  [0.37, 0.42] ^b^** | **21.3  [20.5, 22.0] ^b^** | 85  [82, 88] | **17303  [16223, 18384] ^b^** | **15.4  [14.7, 16.1] ^b^** | **4.29  [3.91, 4.66] ^a^** | 9.12  [8.40, 9.85] | **6.05  [5.94, 6.17] ^b^** | **1.97  [1.78, 2.16] ^b^** | 5.71  [5.26, 6.16] | **3.10  [3.00, 3.20] ^b^** |
|  |  |  | Staygreen | 0.159  [0.140, 0.178] | **0.39  [0.36, 0.42] ^b^** | **22.1  [21.1, 23.1] ^b^** | 87  [82, 91] | **17441  [16002, 18880] ^b^** | **15.3  [14.3, 16.2] ^b^** | **4.07  [3.57, 4.56] ^a^** | 10.13  [9.16, 11.10] | **5.91  [5.76, 6.07] ^b^** | **2.03  [1.74, 2.33] ^b^** | 5.40  [4.80, 6.00] | **3.16  [3.02, 3.30] ^b^** |
|  | **2018** | Early | Non-SG | - | - | 18.8  [18.0, 19.5] | 47  [42, 52] | **12081  [10888, 13274] ^a^** | **18.8  [18.3, 19.4] ^a^** | **3.93  [3.59, 4.26] ^b^** | 8.27  [7.62, 8.93] | 6.34  [6.22, 6.46] | **2.23  [2.06, 2.39] ^a^** | 5.08  [4.66, 5.50] | 3.59  [3.53, 3.65] |
|  |  |  | Staygreen | - | - | **20.1  [19.2, 21.1]*** | 52  [46, 59] | **12660  [11279, 14042] ^a^** | **18.8  [18.1, 19.6] ^a^** | **3.43  [2.95, 3.90] ^b*^** | 8.67  [7.84, 9.51] | 6.24  [6.08, 6.40] | **2.07  [1.84, 2.31] ^a^** | 5.08  [4.54, 5.62] | 3.65  [3.57, 3.73] |
|  |  | Late | Non-SG | 0.159  [0.144, 0.174] | 0.16  [0.14, 0.17] | 19.8  [18.9, 20.6] | 49  [43, 55] | **14986  [13779, 16193] ^b^** | **17.4  [16.8, 18.1] ^b^** | 2.78  [2.39, 3.17] | 8.75  [8.02, 9.49] | 6.02  [5.89, 6.16] | **1.74  [1.55, 1.93] ^b^** | 5.05  [4.58, 5.52] | 3.50  [3.43, 3.57] |
|  |  |  | Staygreen | 0.160  [0.141, 0.178] | 0.16  [0.14, 0.18] | **21.1  [20.2, 22.1]*** | 54  [48, 61] | **15565  [14183, 16947] ^b^** | **17.4  [16.7, 18.2] ^b^** | 3.61  [3.13, 4.08] | 9.15  [8.32, 9.99] | 5.92  [5.77, 6.08] | **2.13  [1.90, 2.37] ^b^** | 5.05  [4.52, 5.59] | 3.56  [3.48, 3.64] |
| **2316b** | **2016** | Early | Non-SG | 0.236  [0.220, 0.253] | **0.42  [0.40, 0.43]^a^** | **18.5  [17.9, 19.1] ^a^** | 56  [53, 58] | 18775  [18019, 19530] | **17.2  [16.7, 17.6] ^a^** | 3.79  [3.12, 4.46] | 9.03  [7.96, 10.10] | **6.28  [6.18, 6.37] ^a^** | 1.88  [1.58, 2.17] | 6.08  [5.59, 6.57] | 3.32  [3.26, 3.38] |
|  |  |  | Staygreen | 0.239  [0.225, 0.253] | **0.40  [0.39, 0.41]^a^** | **18.7  [18.2, 19.2] ^a^** | 56  [54, 58] | 18282  [17650, 18914] | **17.6  [17.2, 17.9] ^a^** | 3.76  [3.2, 4.32] | 9.38  [8.56, 10.21] | **6.32  [6.24, 6.40] ^a^** | 1.82  [1.58, 2.07] | **5.93  [5.52, 6.35] ^a^** | 3.36  [3.31, 3.40] |
|  |  | Late | Non-SG | 0.236  [0.219, 0.252] | **0.37  [0.35, 0.39]^b^** | **20.9  [20.2, 21.5] ^b^** | 55  [53, 58] | 19387  [18630, 20143] | **16.3  [15.8, 16.7] ^b^** | 3.62  [2.95, 4.29] | 9.45  [8.37, 10.52] | **6.06  [5.97, 6.15] ^b^** | 1.69  [1.40, 1.98] | **6.00 [5.51, 6.50] ^a^** | 3.28  [3.22, 3.34] |
|  |  |  | Staygreen | 0.238  [0.224, 0.253] | **0.35  [0.34, 0.37^]b^** | **21.1  [20.5, 21.6] ^b^** | 55  [53, 58] | 18894  [18241, 19547] | **16.6  [16.2, 16.9] ^b^** | 3.58  [3.00, 4.16] | 9.52  [8.65, 10.40] | **6.10  [6.02, 6.18] ^b^** | 1.64  [1.38, 1.89] | **5.85  [5.43, 6.28] ^b^** | 3.32  [3.27, 3.37] |
|  | **2017** | Early | Non-SG | 0.182  [0.167, 0.198] | **0.49  [0.46, 0.51] ^a^** | **18.3  [17.5, 19.1] ^a^** | **97  [93, 100] ^a^** | **16063  [14914, 17212] ^a^** | **17.8  [17.1, 18.6] ^a^** | 4.39  [3.99, 4.79] | 8.68  [7.91, 9.45] | **6.42  [6.30, 6.55] ^a^** | **2.36  [2.15, 2.57] ^a^** | **4.92  [4.44, 5.40] ^b^** | **3.39  [3.28, 3.50] ^a^** |
|  |  |  | Staygreen | 0.182  [0.166, 0.199] | **0.46  [0.43, 0.49] ^a^** | **18.0  [17.1, 18.8] ^a^** | **97  [94, 101] ^a^** | **15010  [13755, 16264] ^a^** | **18.1  [17.3, 18.9] ^a^** | 4.72  [4.29, 5.15] | 8.78  [7.93, 9.62] | **6.46  [6.33, 6.60] ^a^** | **2.21  [1.97, 2.45] ^a^** | 4.91  [4.38, 5.43] | **3.41  [3.29, 3.53] ^a^** |
|  |  | Late | Non-SG | 0.185  [0.168, 0.202] | **0.39  [0.36, 0.42] ^b^** | **21.7  [20.9, 22.6] ^b^** | **82  [78, 86] ^b^** | **19166  [17912, 20421] ^b^** | **15.0  [14.1, 15.8] ^b^** | 4.01  [3.57, 4.44] | 9.85  [9.00, 10.69] | **6.11  [5.97, 6.25] ^b^** | **1.61  [1.37, 1.86] ^b^** | 6.11  [5.59, 6.63] | **3.00  [2.88, 3.12] ^b^** |
|  |  |  | Staygreen | 0.185  [0.170, 0.200] | **0.36  [0.34, 0.39] ^b^** | **21.4  [20.6, 22.2] ^b^** | **83  [79, 86] ^b^** | **18113  [16964, 19262] ^b^** | **15.2  [14.5, 16.0] ^b^** | 4.34  [3.94, 4.73] | 9.94  [9.17, 10.72] | **6.15  [6.03, 6.28] ^b^** | **1.98  [1.77, 2.19] ^b^** | 6.09  [5.62, 6.57] | **3.02  [2.91, 3.13] ^b^** |
|  | **2018** | Early | Non-SG | - | - | 19.0  [18.1, 19.8] | 54  [49, 60] | **12008  [10710, 13306] ^a^** | 18.7  [18.0, 19.3] | **3.76  [3.37, 4.15] ^a^** | 8.83  [8.11, 9.56] | 6.36  [6.22, 6.50] | **2.28  [2.09, 2.47] ^a^** | 4.83  [4.37, 5.30] | 3.55  [3.48, 3.62] |
|  |  |  | Staygreen | - | - | 18.7  [17.8, 19.6] | 54  [47, 60] | **13231  [11751, 14710] ^a^** | 18.6  [17.9, 19.4] | **3.65  [3.17, 4.12] ^a^** | 8.77  [7.94, 9.59] | 6.33  [6.18, 6.49] | **2.12  [1.88, 2.35] ^a^** | 5.08  [4.56, 5.61] | 3.55  [3.47, 3.63] |
|  |  | Late | Non-SG | 0.142  [0.127, 0.157] | 0.14  [0.13, 0.16] | 20.2  [19.4, 21.1] | 53  [47, 58] | **14334  [13122, 15546] ^b^** | 17.8  [17.1, 18.4] | **3.48  [3.1, 3.87] ^b^** | 8.92  [8.19, 9.65] | 6.16  [6.02, 6.29] | **2.09  [1.90, 2.28] ^b^** | 5.24 [4.78, 5.71] | 3.49  [3.42, 3.56] |
|  |  |  | Staygreen | **0.170  [0.155, 0.185]*** | **0.17  [0.16, 0.19]*** | 20.0  [19.2, 20.8] | 52  [46, 58] | **15557  [14345, 16768] ^b^** | 17.7  [17.1, 18.4] | **3.43  [3.04, 3.81] ^b^** | 8.86  [8.13, 9.58] | 6.13  [5.99, 6.27] | **2.03  [1.83, 2.22] ^b^** | 5.49  [5.03, 5.96] | 3.50  [3.43, 3.57] |

Staygreen and Non-SG refers to RILs homozygous for *NAM-A1* (1189a) and *NAM-D1* variants, and those homozygous for the cv. Paragon *NAM-1* allele.
Results of pairwise Tukey *post-hoc* comparisons; **P*<0.05; ***P*<0.01, ****P*<0.001; ^ab^ Significant differences between heading groups (in bold).

**Table S5:** *Differences in grain filling dynamics between ‘early’ heading ± staygreen and parental lines.*

Mean [CI_95%_], results of pairwise Tukey *post-hoc* tests following ANOVA. P-values, * < 0.05, ** < 0.01, *** < 0.001. Corresponding graphs, Figure 9 & Supplemental Figure S3. Grain sampling started at anthesis, which occurred on 15/05/2019 (‘Early’) and 05/06/2019 (‘Late’).

| **Heading Class.** | **Genotype** | **Grain filling component** | **2019 (dd/mm) & days after anthesis (daa)** | | | | | | | | | | |
| --- | --- | --- | --- | --- | --- | --- | --- | --- | --- | --- | --- | --- | --- |
|  |  |  | **12/06  28 daa** | **19/06**  **35 daa** | **24/06**  **40 daa** | **02/07**  **48 daa** | | **10/7**  **56 daa** | **15/07**  **61 daa** | **19/07**  **65 daa** | | **22/07**  **68 daa** | **25/07**  **71 daa** |
| **‘Early’** | **3 x *Ppd-1a* NIL** | **Moisture  content (%)** | 66.3  [64.6, 67.9] | 60.8 [59.1, 62.6] | 54.5 [52.9, 56.2] | 42.2  [40.5, 43.8] | | 39.3 [37.6, 41.1] | 35.8 [34.2, 37.5] | 21.2  [19.6, 22.9] | | 16.4 [14.7, 18.0] | 10.3 [8.6, 11.9] |
|  |  | **Dry grain weight  (10 grains, mg)** | 181 [151, 211] | 214 [183, 246] | 283 [253, 313] | 391 [361, 421] | | 456 [425, 488] | 504 [474, 534] | 509 [479, 539] | | 475 [445, 505] | 546 [516, 576] |
|  | ***Ppd* x 1189a-56** | **Moisture  content (%)** | 67.7 [66.1, 69.4] | 59.7 [58.1, 61.3] | 53.4 [51.7, 55.0] | 46.0 [44.4, 47.7] *** | | 40.6 [39.0, 42.2] | 41.0  [39.2, 42.8] *** | 20.4 [18.5, 22.2] | | 17.7 [16.1, 19.4] | 10.1 [8.44, 11.7] |
|  |  | **Dry grain weight  (10 grains, mg)** | 191 [161, 221] | 272  [242, 302] ** | 321 [291, 351] | 410 [380, 440] | | 487  [457, 517] | 470 [437, 504] | 493 [459, 527] | | 510 [480, 540] | 500 [470, 530] *** |
|  | **3 x *Ppd-1a* NIL** | **Moisture  content (%)** | 69.4 [67.8, 71.1] | 61.0 [59.4, 62.7] | 54.5 [52.8, 56.1] | 46.5  [44.9, 48.2] | | 40.2 [38.6, 41.9] | 35.0 [33.4, 36.7] | 17.2 [15.5, 18.8] | | 16.6 [15.0, 18.3] | 9.7 [8.1, 11.4] |
|  |  | **Dry grain weight  (10 grains, mg)** | 177 [147, 207] | 234 [204, 264] | 319 [289, 349] | 363 [333, 393] | | 515 [485, 545] | 534 [504, 564] | 557  [527, 587] | | 533 [503, 563] | 562 [532, 592] |
|  | ***Ppd* x 2316b-57** | **Moisture  content (%)** | 66.5 [64.9, 68.2] ** | 58.0 [56.3, 59.6] ** | 51.6 [50.0, 53.3] * | 42.6  [40.9, 44.2] *** | | 33.7 [32.1, 35.4] *** | 23.1 [21.5, 24.7] *** | 13.2 [11.5, 15.0] ** | | 14.0 [12.3, 15.6] * | 9.8 [8.2, 11.5] |
|  |  | **Dry grain weight  (10 grains, mg)** | 212 [182, 242] | 293 [263, 323] ** | 373 [343, 403] ** | 397 [367, 427] | | 469 [439, 500] * | 507 [477, 537] | 430  [399, 462] *** | | 460  [430, 490] *** | 467 [437, 497] *** |
| **Heading Class.** | **Genotype** | **Grain filling component** | **2019 (dd/mm) & days after anthesis (daa)** | | | | | | | | | | |
|  |  |  | **01/07**  **26 daa** | | | | **23/07**  **48 daa** | | | | **30/07**  **55 daa** | | |
| **‘Late’** | **Paragon** | **Moisture  content (%)** | 58.5 [56.8, 60.1] | | | | 20.8 [19.1, 22.5] | | | | 16.4  [14.7, 18.1] | | |
|  |  | **Dry grain weight  (10 grains, mg)** | 249  [219, 279] | | | | 418  [386, 450] | | | | 350 [318, 382] | | |
|  | **1189a** | **Moisture  content (%)** | 61.3  [59.5, 63.0] * | | | | 34.6 [32.7, 36.6] *** | | | | 18.1 [16.4, 19.7] | | |
|  |  | **Dry grain weight  (10 grains, mg)** | 214  [183, 246] | | | | 480 [444, 516] ** | | | | 433 [403, 463] *** | | |
|  | **2316b** | **Moisture  content (%)** | 61.5  [59.9, 63.1] ** | | | | 31.7  [30.0, 33.3] *** | | | | 16.8 [15.2, 18.5] | | |
|  |  | **Dry grain weight  (10 grains, mg)** | 231  [201, 261] | | | | 449 [419, 479] | | | | 384 [354, 414] | | |

**Table S6:** *NAM-1 mutations are associated with grain protein content reduction*.

|  | | | | **2016** | | | **2017** | | | **2018** | | |
| --- | --- | --- | --- | --- | --- | --- | --- | --- | --- | --- | --- | --- |
| **Mutant** | **RILs** | **Site** | **Heading** | **Staygreen** | **Non-SG** | **% Difference (SG vs. Non-SG)** | **Staygreen** | **Non-SG** | **% Difference (SG vs. Non-SG)** | **Staygreen** | **Non-SG** | **% Difference (SG vs. Non-SG)** |
| **1189a (*NAM-A1*)** | *Ppd* x staygreen | Cambs. | ‘Early’ | **13.0 ± 0.2** | **14.1 ± 0.3** | **-7.8% ***** | **12.6 ± 0.3** | **13.6 ± 0.2** | **-7.4% *** | 12.0 ± 0.4 | 12.2 ± 0.3 | -1.6% |
|  |  |  | ‘Late’ | **13.3 ± 0.2** | **14.4 ± 0.2** | **-7.6% **** | **13.4 ± 0.3** | **14.4 ± 0.2** | **-6.9% *** | 11.5 ± 0.4 | 11.7 ± 0.4 | -1.7% |
|  |  |  | Overall | **13.2 ± 0.2** | **14.2 ± 0.2** | **-7.0% ***** | **13.0 ± 0.3** | **14.0 ± 0.2** | **-7.1% **** | 11.7 ± 0.4 | 12.0 ± 0.3 | -2.5% |
|  |  | France | ‘Early’ | - | - | - | 12.0 ± 0.4 | 12 ± 0.3 | - | **11.9 ± 0.2** | **13.1 ± 0.2** | **-9.2% **** |
|  |  | Germany | ‘Early’ | - | - | - | - | - | - | **13.4 ± 0.2** | **14.1 ± 0.2** | **-5.0% *** |
|  | Paragon x staygreen | Norwich | ‘Late’ | - | - | - | **11.9 ± 0.1** | **12.4 ± 0.1** | **-4.0% ***** | **11.3 ± 0.0** | **12.0 ± 0.1** | **-5.8% ***** |
| **2316b (*NAM-D1*)** | *Ppd* x staygreen | Cambs. | ‘Early’ | **13.7 ± 0.2** | **14.6 ± 0.2** | **-6.2% **** | 13.3 ± 0.3 | 13.8 ± 0.2 | -3.6% | 11.6 ± 0.4 | 12.2 ± 0.4 | -4.9% |
|  |  |  | ‘Late’ | **13.9 ± 0.2** | **14.8 ± 0.2** | **-6.1% **** | 14.9 ± 0.2 | 15.4 ± 0.3 | -3.2% | 11.8 ± 0.4 | 12.3 ± 0.4 | -4.1% |
|  |  |  | Overall | **13.8 ± 0.2** | **14.7 ± 0.2** | **-6.1% **** | 14.1 ± 0.2 | 14.6 ± 0.2 | -3.4% | 11.7 ± 0.3 | 12.3 ± 0.3 | -4.9% |
|  |  | France | ‘Early’ | - | - | - | 12.4 ± 0.4 | 11.9 ± 0.3 | +4.2% | 12.7 ± 0.2 | 12.9 ± 0.2 | -1.6% |
|  |  | Germany | ‘Early’ | - | - | - | - | - | - | 13.5 ± 0.2 | 14.2 ± 0.2 | -4.9% |
|  | Paragon x staygreen | Norwich | ‘Late’ | - | - | - | **11.7 ± 0.1** | **12.1 ± 0.1** | **-3.3% **** | **11.8 ± 0.1** | **12.2 ± 0.1** | **-3.2% ***** |

Grain protein content (%) ± SE
Results of pairwise Tukey *post-hoc* comparisons; **P*<0.05; ***P*<0.01, ****P*<0.001 (in bold)
